# Structural and dynamical basis for the interaction of HSP70-EEVD with JDP Sis1

**DOI:** 10.1101/2022.11.26.517237

**Authors:** Carolina O. Matos, Glaucia M.S. Pinheiro, Icaro P. Caruso, Gisele C. Amorim, Fabio C. L. Almeida, Carlos H. I. Ramos

**Author notes:** Correspondence should be addressed to FCLA, and CHR.

## Abstract

We employed NMR spectroscopy to investigate the structure and dynamics of the class B J domain protein (JDP) of S. cerevisiae (Sis1) complexed with an EEVD peptide of HSP70. It is widely recognized that the interactions between the EEVD motif and Sis1 play a crucial role in the chaperone activity. Notably, the deletion of the EEVD impairs the ability of Sis1 to bind with HSP70, while leaving the interaction between the class A JDP Ydj1 and HSP70 unaffected. Leveraging the advantages of NMR, which is particularly suitable for studying transient interactions, we provide compelling evidence that the EEVD motif transiently engages multiple sites on Sis1. Our findings revealed that EEVD binds to two distinct sites within the C-terminal domain I (CTDI) of Sis1. The interaction at these sites plays a crucial role in anchoring HSP70 to Sis1 at site I, as well as displacing the client protein at site II. Notably, site II is also the binding site for the client protein, and its displacement occurs through competition with the binding to site II. In addition to these interactions, we observed that EEVD, as a transient electrostatic binder, also interacts with the J domain and the GF-rich loop located between the J domain and α-helix 6. We propose that the interaction between EEVD and Sis1 facilitates the dissociation of α-helix 6, promoting a conformational state that is more favorable for interaction with HSP70 at the nucleotide-binding domain (NBD) and substrate-binding domain (SBD) interface. We also employed α-synuclein as a substrate to investigate the competitive nature between EEVD and the client protein. Our experimental findings provided evidence supporting the interaction of EEVD with the client protein at multiple sites. Our findings contribute essential insights into the mechanistic cycle of class B JDPs, paving the way toward a more complete understanding of the primary function of Sis1, which is the transfer of the client protein to HSP70, where multiple site transient interactions play a collective role.

## Introduction

Chaperones of the 70 kDa heat shock protein (HSP70) superfamily are key components of the cellular proteostasis system which are composed of a nucleotide-binding domain (NBD), a substrate binding domain (SBD), and a C-terminal disordered region ending in an EEVD motif ^1–3^. The SBD is composed of the SBDβ subdomain, also known as the base subdomain, and the SBDα subdomain, the lid subdomain. The EEVD motif is involved in the intramolecular regulation of HSP70 function and intermolecular interactions with J-domain proteins (JDPs, also known as HSP40s or DNAJs), a family of cochaperones involved in the delivery of client-proteins. JDPs bind to the interface between the NBD and the SBD of HSP70 and it is necessary for stimulation of its ATPase activity^4,5^.

JDPs are classified into three classes, which are all characterized by the presence of a J-domain, N-terminal in classes A and B, and anywhere in class C. A disordered region follows the J-domain, which is composed of a glycine-phenylalanine-rich (GF) in class A and a GF region followed by a glycine/methionine-rich (GM) in class B. Both classes A and B have homologous C-terminal β-barrel domains (divided into CTDI and CTDII) and a dimerization domain (DD). The CTDI of class A contains a zinc-finger-like region (ZFLR) which is not present in class B (Fig. 1)^6–9^.

**Fig. 1.**
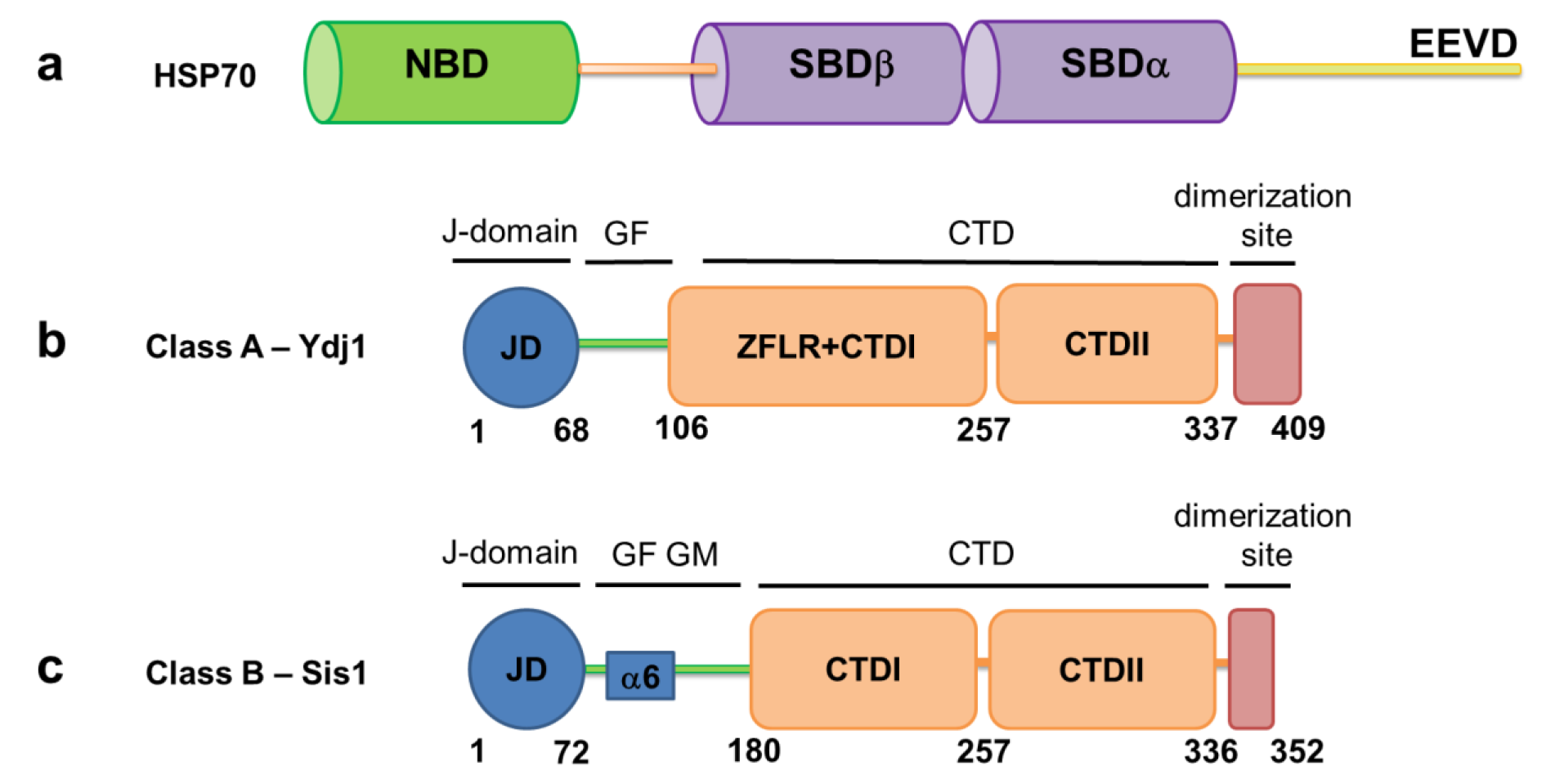
Domain architecture of HSP70 and JDPs. **a** Schematic representation of HSP70 chaperones, composed of a nucleotide-binding domain (NBD, which binds ATP) in green, substrate binding domain (SBD) in purple, divided into SBDβ (base) and SBDα (lid) subdomains, that are connected by a flexible linker (orange) and a C-terminal disordered region terminating in a high conserved EEVD motif (yellow).**b** Schematic representation of the domain organization of Ydj1, class A, and **c** Sis1, class, B from yeast. The different domains are marked as follows: JD: J-domain (blue, contains α-helices 1-5); GF: Gly-Phe rich region and GM: Gly-Met rich region (green). The GF region of Sis1 contains the α-helix 6, which is conserved in class B JDPs and is described as α-helix 5 for DNAJB1^15^ and others; ZFLR: zinc-finger like region; CTD: C-terminal domain (I and II; orange) and dimerization domain (DD, red).

Two cytosolic and dimeric JDPs from *Saccharomyces cerevisiae*, the class A Ydj1 and the class B Sis1, stimulate the ATPase activity of HSP70 and its function^10^. Remarkably, *sis1* is an essential gene while *ydj1* is not. Cells knocked out of the *ydj1* gene show a phenotype of severe growth defect and a decrease in stress tolerance, while the overexpression of Sis1 can partially suppress its slow growth phenotype. Nevertheless, overexpression of Ydj1 does not restore the viability of cells knocked out of the *sis1* gene^11,12^. It is well-established that the EEVD motif of yeast HSP70/Ssa1 can bind to Sis1 by electrostatic interactions, which is functionally important for its chaperone activity^13^. Deletion of the EEVD (HSP70_ΔEEVD_) disrupts the ability of Sis1 to associate with HSP70 but does not affect the interaction between Ydj1 and HSP70. Substitution of the J-domain of Ydj1 or Xdj1 (a paralog of Ydj1) for that of Sis1, results in the modified Sis1 associating with HSP70_ΔEEVD_. Sis1-EEVD interaction is also important for Ssa1 (*S. cerevisiae* HSP70) recruitment to bind in aggregated substrates^14^.

An autoinhibitory mechanism for HSP70 binding was shown for the class B JDPs DNAJB1 and DNAJB6^15,16^, in which the binding site located at the J-domain is occluded by a conserved α-helix 5 at the GF region (Fig. 1c). The occlusion can be reverted by the interaction with the HSP70-EEVD motif. This self-regulation is unique for class B JDPs and is essential for the disaggregation of amyloid fibers by HSP70/DNAJB1^15^. Our group recently showed the key importance of the transient interaction of Sis1 to HSP70^17^. Sis1 J-domain binds to HSP70 through a hydrophobic and positively charged patch mainly at α-helices 2 and 3 with the participation of the conserved HPD motif, located between these helices, and is released by the addition of ATP^17^. In the context of the full-length Sis1, the J-domain is in a transient intermediate conformation, in which the HSP70- interacting patch is protected by internal transient interactions^17^.

Structural information on the interaction between JDPs and the EEVD motif is still scarce, and there are many open questions about the biological effects caused by the association of these chaperones. A better understanding of the EEVD interaction with JDP arises from the results of this work, which gives a detailed NMR characterization of the dynamics and interaction between full-length Sis1 (residues 1-352, herein referred to as Sis1_1-352_) and the HSP70-EEVD motif. Characterization of the dynamics and interaction between the Sis1 J-domain (residues 1-81, herein referred to as Sis1_1-81_) and the HSP70-EEVD motif was also performed to reveal more details about the interaction. We proposed models taking into consideration the competing and simultaneous binding of the client protein and EEVD to the globular domain CTDI and the intrinsically disordered region GF/GM, which may generate entropic forces to orient the transfer of the client protein to HSP70.

## Results

### The interdomain dynamics of Sis1_1-352_ is affected by EEVD

The understanding of the interaction between the HSP70 C-terminus EEVD motif with the cochaperone Sis1 is important because the deletion of this motif abolishes the ability of Sis1 to partner with HSP70 and pursue its chaperone activity^14^. The results were based on 70% of the backbone resonance assignment of the full-length protein, Sis1_1-352_ (BMRB ID 51817). We first studied the dynamics of free Sis1 and the changes elicited by the interaction of a GPTIEEVD peptide (representing the EEVD motif). The ^15^N longitudinal (R_1_) and transverse (R_2_) relaxation rates of Sis1_1-352_ free and bound to the EEVD peptide were measured. Table 1 describes the ^15^N R_2_/R_1_ ratio and the apparent rotational diffusion (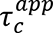) of each domain in the context of the full-length protein^18^. 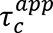 reflects the rotational freedom of each domain in the free and EEVD-bound states and was calculated by filtering out residues in conformational exchange or thermal flexibility using the average ^15^N R_2_/R_1_ of each region considering only the values within the average plus or minus one standard deviation (sd).

**Table 1.** ^15^N longitudinal (R_1_) and transverse (R_2_) relaxation rates (R_2_/R_1_) and the apparent rotational diffusion (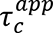) of free and EEVD-bound Sis1_1-352_

| Domain (free or bound) | $^{15}\text{N-R}_2/\text{R}_1$ | $\tau_c^{app}, \text{ s}$ |
| --- | --- | --- |
| J-domain (free) | $22 \pm 2$ | $9.8 \pm 1.0 (\times 10^{-9})^*$ |
| J-domain (bound) | $20 \pm 2$ | $9.2 \pm 1.0 (\times 10^{-9})^*$ |
| GF/GM region (free) | $30 \pm 13$ | $11 \pm 5 (\times 10^{-9})$ |
| GF/GM region (bound) | $22 \pm 8$ | $10 \pm 4 (\times 10^{-9})$ |
| CTD (free) | $70 \pm 8$ | $18 \pm 2 (\times 10^{-9})$ |
| CTD (bound) | $71 \pm 8$ | $18 \pm 2 (\times 10^{-9})$ |
| * $\tau_c^{app}$ of the bound domain is lower than that of a free domain at a significance level of $P < 0.05$ | | |

The relaxation parameters for free Sis1_1-352_ (Fig. 2a, Table 1) highlighted the differences in dynamics between its extended J-domain (1-72), GF (73-121), GM (122- 178), CTDI (179-257), CTD II (258-335) and dimerization (DD, 336-352) domains. We measured a significant difference of ^15^N R_2_/R_1_ for the J-domain (22 ± 2), GF/GM (30 ± 13), and CTD (70 ± 8), indicating that the domains located at the C-terminus, CTD, and DD, have the most restricted rotational freedom. The calculated apparent rotational correlation time (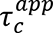) showed that all domains had more rotational freedom than expected for a rigid globular 75.2 kDa protein, considering the scissor-like anisotropic structure of Sis1. The CTD showed the most restricted domain motion (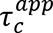 = 18 ± 2 ns), possibly due to its proximity to the dimerization domain. The J-domain and the GF/GM regions behaved as a highly flexible arm with 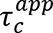 of 9.8 ± 1.0 ns and 11.0 ± 5.0 ns, respectively. Note that the GF/GM region (sd=5) presents a higher dispersion of ^15^N R_2_/R_1_ values when compared to the J domain (sd = 1) or CTD (sd = 2).

**Fig. 2.**
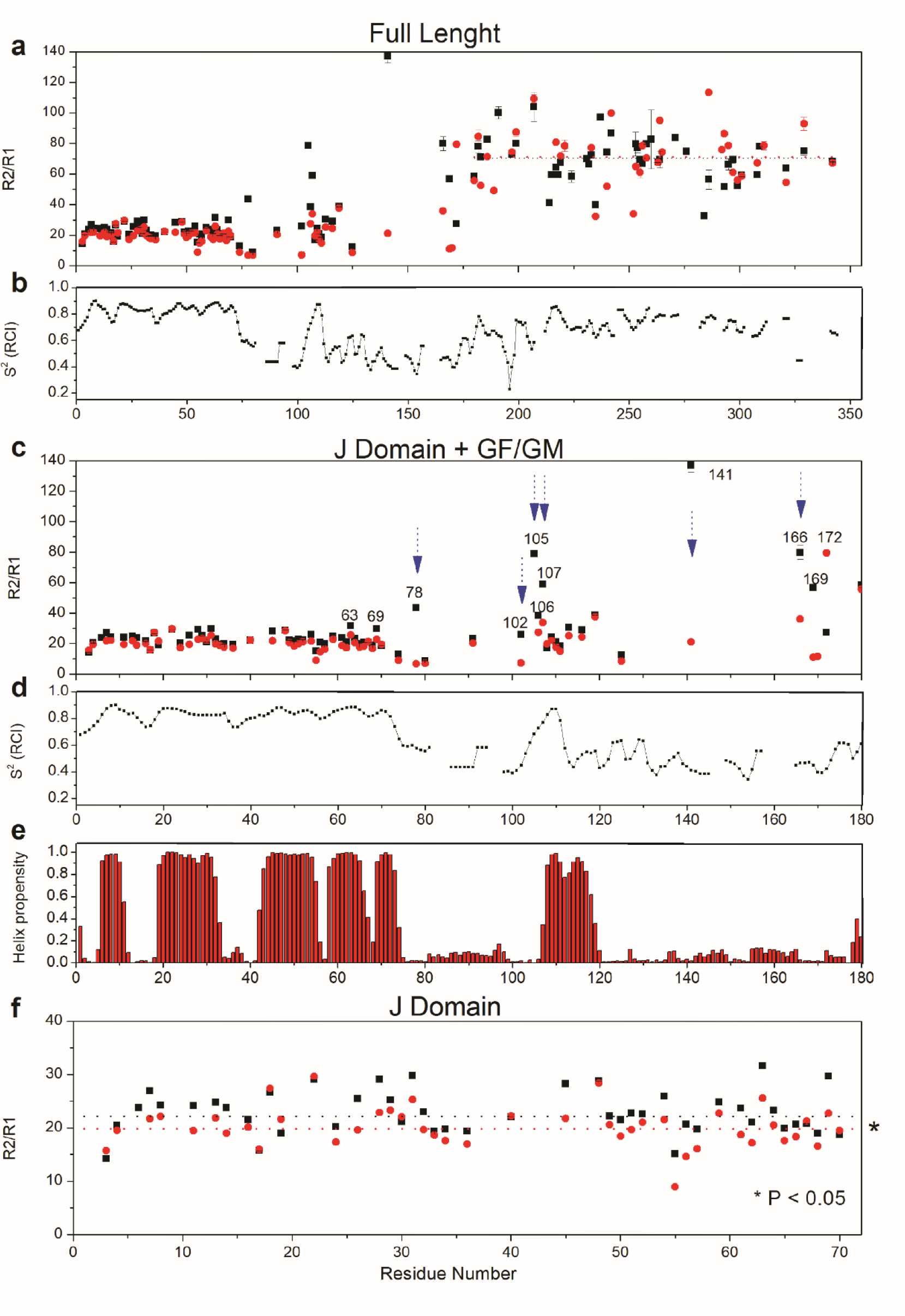
Relaxation parameters, chemical shift-based order parameter, and Talos-N secondary structure of Sis1_1-352_. **a** ^15^N-R_2_/R_1_ for Sis1_1-352_ (full-length) free (black squares) and bound to EEVD (red circles) for each residue. The dotted lines show the average of ^15^N-R_2_/R_1_ for the CTDI, CTDII, and dimerization domains. **b** Chemical shift-based order parameter (S^2^) for residues of free Sis1_1-352_. The lower the value of S^2^ the higher the internal flexibility. **c** Zoomed plot (residues 1 to 180) showing ^15^N-R_2_/R_1_ for J-domain and GF/GM region for the free (black squares) and EEVD-bound state of Sis1_1-352_ (red circles). Blue arrows highlight the residues for which ^15^N-R_2_/R_1_ decreased upon binding to EEVD. **d** Zoomed plot (residues 1 to 180) showing S^2^ for J-domain and GF/GM region for the free Sis1_1-352_. **e** Zoomed plot (residues 1 to 180) showing the helical propensity predicted by Talos-N for J-domain and GF/GM region for the free Sis1_1-352_. **f** Zoomed plot (residues 1 to 72) showing ^15^N-R_2_/R_1_ for the J-domain free (black squares) and EEVD-bound (red circles). Dotted lines, average of ^15^N-R_2_/R_1_. *, statistically significant for P < 0.05 decrease in average ^15^N-R_2_/R_1_.

The chemical shift-based order parameter (S^2^) analysis, for Sis1_1-352_ (Fig. 2b) and Sis1_1-180_ (Fig. 2d), was obtained using TALOS-N^19,20^ and was informative on the structural order of each of the regions/domains of the protein. We observed high order for the well-folded J-domain, CTDI, CTDII, and dimerization domain (DD). On the other hand, GF/GM showed high backbone flexibility, typical of an IDR, except for the residues 104 to 112, which are highly ordered. The region 104-112 contains the N-terminal portion of α-helix 6, which spans from 107 to 119 (Fig. 2e). Although the higher order for the helical segment comes as no surprise, remarkably, there was a mismatch between the ordered region within the GF (104-112, Fig 2d) and α-helix 6 (107-119, Fig 2e).

The orthologous human DNAJB1 also shows an α-helix (α-helix 5) within the GF region that blocks the J-domain interacting site to HSP70, as an autoinhibitory mechanism^15^. A similar GF α-helix (named α-helix 5) is present in the NMR structure of DNAJB6^16^. The authors also suggest it competes with DnaK/HSP70 for binding to the J-domain. Sis1 structural predictions made by Alphafold^21,22^ also suggest the interaction of α-helix 6 with the J-domain, indicating that Sis1 is also regulated by an autoinhibitory mechanism. The observed order in this region (residues 104-112, Fig. 2b) supports this interaction, nevertheless, the observed mismatch between the ordered (104-112) and helical residues (107-119), suggests an on-off equilibrium between a J-domain-associated (on-state) and a free, or J-domain-dissociated (off-state) α-helix 6. In this situation, the assigned chemical shift in solution reflects the average between the J-associated and J-dissociated α-helix 6, similar to what was observed for DNAJB1^15^ and DNAJB6^16^. Furthermore, the lower structural order observed for the C-terminal portion of the α-helix 6 (which is positioned between residues 113-119) could be attributed to a partial or total unfolding of the helix in the off-state. This agrees with secondary structure analyses by Jpred4^23^, which does not predict a well-folded α-helix 6. Jpred4 informs on secondary structure tendency, based solely on the sequence, meaning that the off state (J-domain-dissociated) would be disordered. Another evidence of an on-off equilibrium is that residues S78, G102, A105, F106, and S107, which are in the GF region and N-terminal to α-helix 6, are in conformational exchange, indicated by their high ^15^N R_2_/R_1_ (Fig. 2c).

We evaluated the effect of the EEVD-peptide binding on the dynamics of Sis1_1- 352_ (Fig. 2). The presence of the EEVD led to a subtle but significant increase in the rotational freedom of the J-domain. 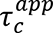 decreased from 9.8 ± 1.0 ns to 9.2 ± 1.0 ns, which is statistically significant (P < 0.05, Fig. 2f, Table 1), given the high number of measurements (35 measurements of ^15^N R_2_/R_1_ in the absence and 37 in the presence of the EEVD). Note that most of the ^15^N R_2_/R_1_ in the presence of EEVD had a decrease in value in Fig. 2f.

The decrease in ^15^N R_2_/R_1_ for the residues S78, G102, A105, F106, and S107 (Figs. 2a and 2c) suggested that the on-off equilibrium of α-helix 6 was suppressed by the presence of the EEVD. We also observed a significant decrease in ^15^N R_2_/R_1_ for the residues G141, G166, and S169 (Figs. 2a and 2c), suggesting a quenching in the conformational exchange of residues that are at the GM region. Residue S172 was the only one with increased ^15^N R_2_/R_1_ elicited by the binding of EEVD. However, a significant change in 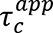 of the CTD was not observed when EEVD was present. In conclusion, altogether, the results indicated significant changes in the dynamics of *Sis1*_1- 352_ caused by the binding of EEVD.

### EEVD interacts in multiple sites within Sis1_1-352_

To further understand the effect that the EEVD binding causes in Sis1_1-352_ we used two methods to map the binding sites. Paramagnetic relaxation enhancement (PRE) on Sis1_1-352_ residues elicited by an N-terminal TEMPO-labeled EEVD peptide (Fig. 3a) and the chemical shift perturbation (CSP) caused by the binding to the EEVD peptide (Fig. 3b, Fig. S1). We assigned 70% of the backbone of Sis1_1-352_ and, to have more reliability in the conclusions, all the overlapping peaks from the analysis were removed. We mapped two major regions of interaction, guided by the residues presenting PRE < 0.88 (1-182) or 0.7 (183-352) (dotted lines in Fig 3a) or CSP > 0.018 (dotted line in Fig 3b): (i) the GF IDR, spanning from residue 78 to 102, which is between the J-domain and α-helix 6, and (ii) part of GM and the CTDI domain (residues 180-256). Interestingly, both regions in free Sis1_1-352_ contain residues in conformational exchange, as shown by their above-the-average values of ^15^N-R_2_/R_1_ (Fig. 2a). In the presence of EEVD the ^15^N-R_2_/R_1_ dropped to average values, meaning the milli-to microsecond motions (conformational exchange) were suppressed by the binding.

**Fig. 3.**
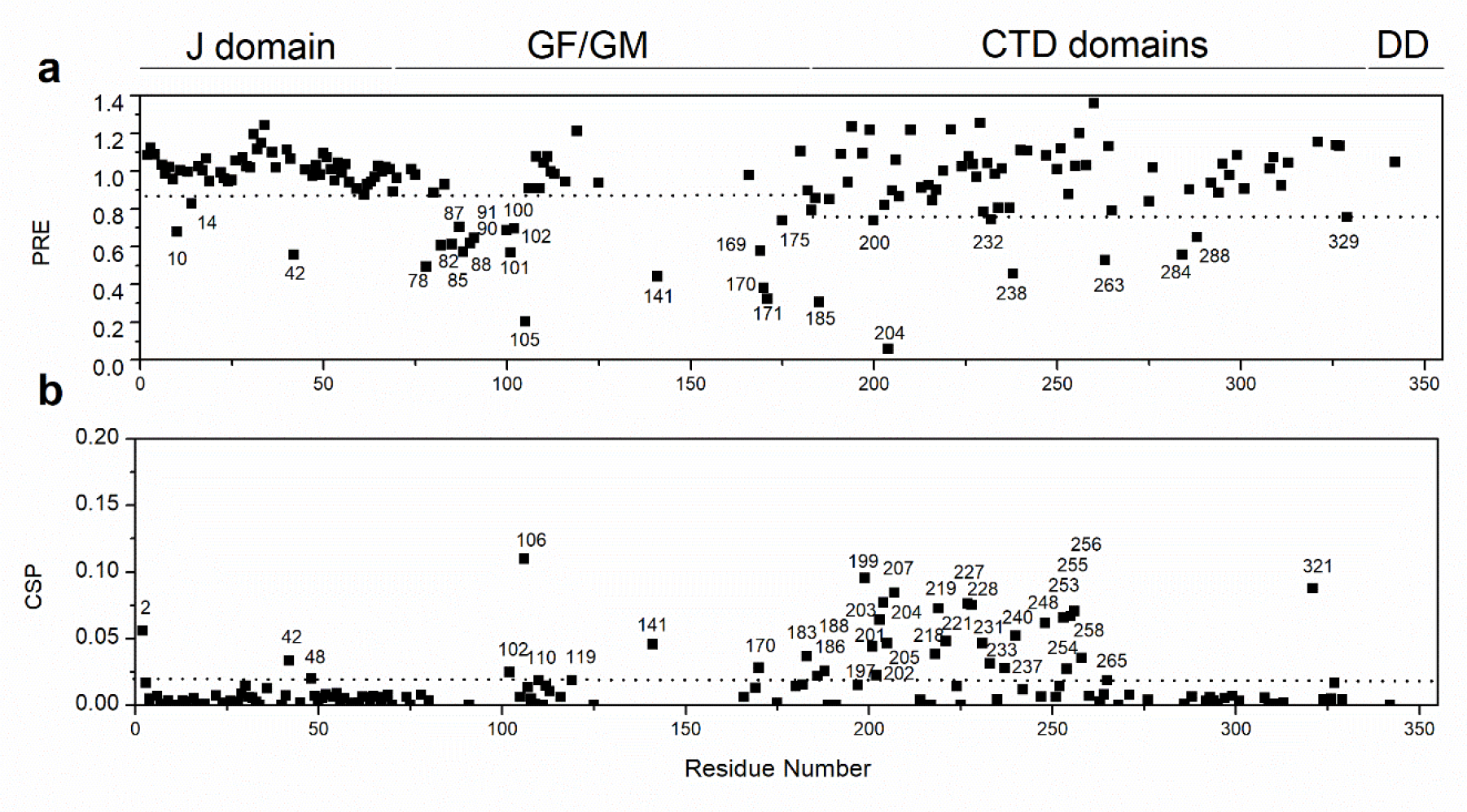
EEVD binding to Sis1_1-352_. **a** Experimental intensity ratios of backbone amide Sis1_1-352_ (full-length) in complex with an EEVD paramagnetic peptide (see Material and Methods) for each residue. An intensity ratio of 1 indicates no effect of the spin label on an amide proton. Residues with a significant PRE are indicated. PRE is the normalized intensity ratio (*I^para^*/*I^dia^*) between the paramagnetic and diamagnetic (reduced with ascorbate) TROSY spectra. **b** Chemical shift perturbation of Sis1_1-352_ resonances upon the addition of the EEVD peptide. CSPs larger than the mean plus one standard deviation are indicated by dotted lines.

The observed PRE and CSP in multiple regions of the protein indicated that the EEVD binds to more than one site (majorly GF and CTDI). Supporting this hypothesis was the observation of PRE and CSP for residues in the J-domain, GF, and CTDI, and also few residues at the GM region just before CTDI (Fig. 3). CSP measures the change in the chemical environment elicited by the presence of the ligand what could result from direct or indirect contact with the ligand. PRE, on the other hand, is a direct distance measurement of the ligand to the protein. PRE broadens the line of any nuclear spin that is less than ∼30 Å of the paramagnetic center^24^. PRE effect was measured by the significant decrease of *I^para^*/*I^dia^*, which demonstrated the direct interaction of EEVD in more than one site, corroborating the CSP measurements. For the J domain, we observed both CSP and PRE: V2 (CSP), A10 (PRE), S14 (PRE), T42 (PRE and CSP), I48 (CSP). For de GF domain we also observed CSP and PRE: S78, G82, G85, A87, G88, G90, A91, F100, S101, G102, and A105 (CSP) and G102, F106, D110, and F119 (PRE). Interestingly, PRE effect was observed on the N-terminal vicinity of α-helix 6 (G102 and F106) and at α-helix 6 (D110, exposed to the solvent, and F119, facing the J domain interacting patch, α2-loop-α3). For CTDI and the GM region, just before CTDI, we observed many residues with CSP and PRE. We observed PRE for the following residues: S169, A170, S171, and T175 are at the GM pre-CTDI, and N185, S200, G204, I232, Q238, at CTDI. We observed CSP for the following residues: A170 (GM, pre-CTDI), Q183, L186, V188, K197, K199, F201, K202, I203, G204, R205, D218, I219, L221, A227, G228, I231, K233, G237, D238, N240, T248, I253, Q254, E255, K256, H258 and G265 (CTDI). Finally, NMR data-derived docking of the EEVD peptide to Sis1_1-352_ supported the multiple binding sites hypothesis.

### Deriving structural models for the interaction of EEVD with Sis1_1-352_

Haddock^25^ was used to dock the EEVD peptide into the different regions of Sis1_1-352_. First, the docking targeted the dimeric CTDI (180-257) using only the PRE and CSP data relative to this region. We observed poses of the EEVD-peptide binding to the CTDI in mainly two interaction sites, named here I and II, which are in opposite faces of the domain (Figs. 4 and S2). The interaction space at the CTDI (sites I and II) was well-defined, but not the orientation and position of the EEVD-peptide relative to these sites because it, allowing for different poses. The strategy for analysis focused on describing the interaction space within the CTDI and was based on the many observed poses of the EEVD relative to this domain (Fig. S2c). For that, we selected the 100 lowest binding energy conformers, which varied from -399.8 to -174.0 kcal/mol (Fig. S2a). Similar results would be also obtained if we had chosen the 100 lowest energy structures because of the linear correlation observed between the energy of the complex and binding energy (Fig. S2b). Note that by using the 100 lowest binding energy structures we warrant that all selected structures had negative complex and binding energy (Fig. S2b), representing well-behaved complex models in terms of geometry.

**Fig. 4.**
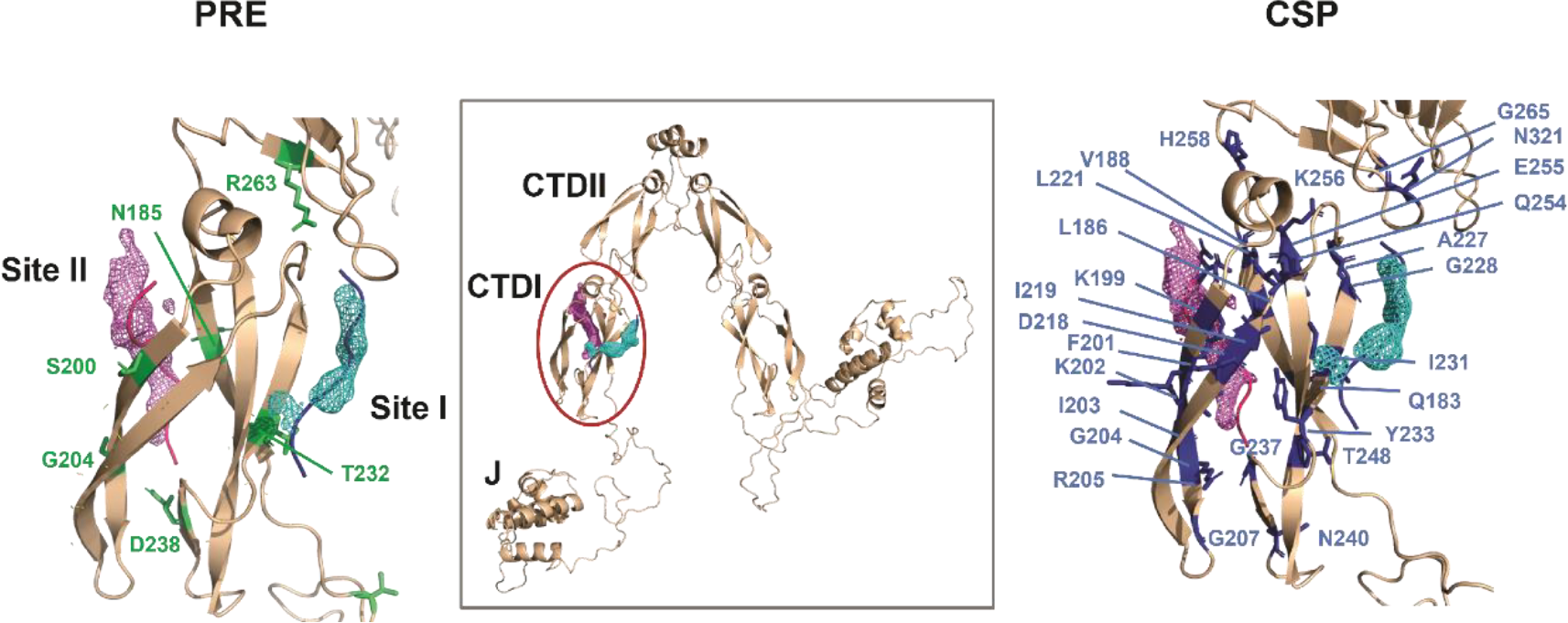
EEVD binds to multiple sites at CTDI of Sis1_1-352_. Cartoon representation of the Sis1 structure predicted by AlphaFold highlighting the CTDI, in which the atomic probability density map of the EEVD peptide at sites I (cyan) and II (magenta) are shown. The residues mapped by PRE (left) and CSP (right) are labeled in green and blue, respectively. Site II is formed by the motif β1, α, β2, while site I is by β4.

**Fig. 5.**
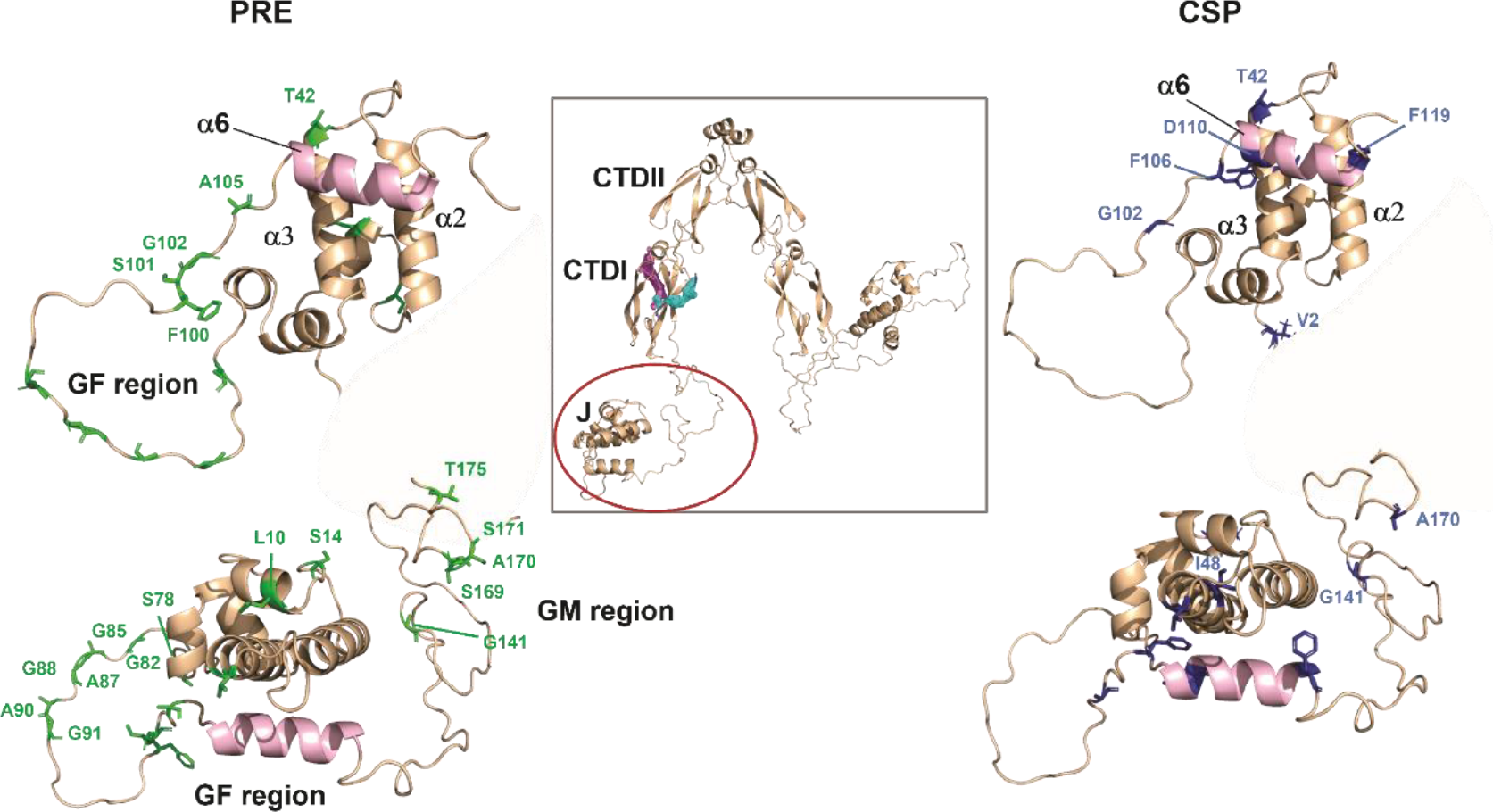
EEVD binds to multiple sites at J domain, GF, and GM region of Sis1_1-352_. Cartoon representation of the Sis1 structure predicted by AlphaFold highlighting the J domain and the GF region. The residues mapped by PRE (left) and CSP (right) are labeled in green and blue, respectively. The α-helix 6 is highlighted in green and was predicted to be in the autoinhibited conformation, blocking the J domain binding patch^15,17^. Note that the residues T42 and I48, at α- helix 3, display either PRE or CSP suggesting the involvement of the J domain in the interaction with EEVD.

Next, we separated the 100 structures into two clusters: cluster I, containing all the poses that bind to site I (73 structures), and cluster II containing the poses that bind to site II (27 structures) (Fig. S2c). Using *Volmap 1.9.3*, a structural tool of VMD software^26^, we calculated the atomistic (mass) probability density map for both clusters. The atomic probability density map for site I (cluster I) and site II (cluster II) are described by the meshes in cyan (site I) and magenta (site II) in Fig. 4. Note that the position of the density maps matches with the experimental PRE (Fig. 4a) and CSP (Fig. 4b) in the CTDI as measured for Sis1_1-352_ in the presence of the EEVD peptide. Figs. S2d and S2e show the lowest binding energy poses, which superpose well with the calculated density maps. For site I, the density map also coincides exactly with the position of an HSP70-EEVD peptide complexed with the human DNAJB1^27^ (Fig. S3a). One of the poses even forms an antiparallel β-strand with the CTDI β-strand 4 (β4), the same configuration of the EEVD complex at site I of human DNAB1^27^. For site II, the density map is between β1 and β2, while both crystal structures describe the interaction of EEVD peptide with either DNAB1 (3AGY, Fig. S3b) or Sis1 (2B26, Fig. S3c) showing interaction with β2, forming and antiparallel β-strand. To assert the stability of the docking presented here, we run 100 ns molecular dynamics (MD) simulations of the 2 lowest energy docking poses for each site (I and II). As a control, we also run the MD simulations starting from the crystallographic structure of EEVD complexed with DNAJB1 and Sis1, at sites I and II (Fig. S4 and S5). The docked structures were stable along the MD simulations.

At this point, for the sake of comparison, it is important to analyze the electron density maps of all available crystal structures describing the interaction of the EEVD peptide and the CTDI. In a crystal structure of Sis1_171–352_ complexed with the HSP70- GPTVEEVD peptide^13^ (PDB 2B26), the EEVD backbone, which is bound to site II, is well determined but none of the side chains has a well-defined electron density, making it difficult to be assertive on the orientation of the peptide relative to β2 (see Fig. S6a and S6B). It is also important to mention that, in this case, the EEVD peptide is not involved in any crystal contact.

In a crystal structure of DNAJB1^27^ complexed with the EEVD peptide (PDB 3AGY), at site II, the same observations are valid, with low electron density for the EEVD-peptide backbone and none for the side chains (Fig. S6c and S6d). For DNAJB1^27^ at site I, the electron density for the backbone is well-defined and sidechains P2, I4, E5, and E6 have reasonably well-defined electron densities, enabling the assertiveness of the relative orientation of the EEVD-peptide and β4, as antiparallel β-strands (Fig. S6c and S6e).

The analysis of the lowest energy docking models generated in this work for site I from Sis1, in which the atomic probability map coincides with the electron density map of the crystal structures, allows for two possible orientations of the peptide relative to β4 (Fig. S7). The EEVD-peptide lowest energy structure (Fig. S7a) has a parallel orientation relative to the CTDI-β4, while the 2^nd^ lowest energy structure has an antiparallel orientation to this domain (Fig. S7b), similar to the EEVD peptide in DNAJB1 at site I (Fig. S7c). Remarkably, both orientations are stabilized by a considerable number of salt bridges, hydrophobic and hydrogen bond interactions (Fig. S8 and Table 2).

**Table 2.** Orientation and salt bridges from the CTDI:EEVD-peptide complexes at sites I and II

| Complex | Orientation | Salt-bridges |
| --- | --- | --- |
| Site I |  |  |
| Lowest energy (Sis1, this work) | Parallel to $\beta 4$ | K234/D8, K230/E5 |
| 2nd Lowest energy (Sis1, this work) | Antiparallel to $\beta 4$ | K230/E6, K222/E5, K226/D8 |
| DNAJB1<sup>27</sup> (crystal structure, 3AGY) | Antiparallel to $\beta 4^*$ | K217/E5 and K213/D8 (if D8 were present) |
| Site II |  |  |
| Lowest energy (Sis1, this work) | Parallel to $\beta 1$ and antiparallel to $\beta 2$ | K199/E6, K199/D8, K256/D8 |
| 2nd Lowest energy (Sis1, this work) | Parallel to $\beta 1$ and antiparallel to $\beta 2$ | K256/E5, K197/D8 |
| DNAJB1<sup>27</sup> (crystal structure, 3AGY) | Antiparallel to $\beta 2^{**}$ | K181/E6, K184/E5, K182/D8 |
| Sis1<sup>13</sup> (crystal structure, 2B26) | Antiparallel to $\beta 2^{**}$ | K214/E6, K202/E6, K199/D8 |

\* Electron density enables assertiveness on the orientation.
\*\* Electron density does not enable assertiveness on the orientation

Similar analyses were performed for site II and the atomic probability density maps of the EEVD-peptide (Fig. S3b and S3c) did not coincide with those of the electron densities for the crystal structures of the crystal complexes of either Sis1^13^ or DNAJB1^27^. The position and orientation resulting from the docking models suggest that the EEVD-peptide binds between β1 and β2 of CTDI, which is compatible with the CSP and PRE data (Fig. 4). The relative orientation to β2 of the lowest energy complexes is the same as those depicted in the crystal structures (antiparallel to β2, Fig. S7d-g). The lowest energy structures are stabilized by a considerable number of salt bridges (Table 2, Fig. S9), hydrogen bonds, and hydrophobic interactions.

The interaction of the J-domain with the EEVD-peptide is largely electrostatic, being stabilized by 2 salt bridges. The interaction of J-domain with HSP70 forms 5 salt bridges: R27/D211, D35/R167, R22/E206 and K26/E217, with the NBD and K48/D477 with the SBDβ domains^28^, i.e. it is also largely electrostatic. EEVD peptide is negatively charged, being able to compete with the salt bridges stabilizing the J-domain/HSP70 interaction. It is important to mention that the only available structure of the J-domain complexed with HSP70 is that of the *E.coli* DNAJ^28^, nevertheless, all the charged residues (R22, K26, R27, D35, and K48) involved in intermolecular salt-bridges with DNAJ are conserved in the J-domain of Sis1 (K23, R27, K28, D36, and K46, respectively)

### Binding to a client protein and EEVD competition

We asked whether the EEVD peptide competes for the binding site of a client protein since from this result it is possible to propose some more important steps to the DNAJ-HSP70 interaction model. For that, we used α-synuclein (α-syn), a well-known interactor of chaperones^29–31^, as a client protein. We added 2:1 stoichiometry of α-syn to Sis1_1-352_ and measured the resultant CSP (Fig. 6, S10). We observed the most significant CSPs for residues of GF region (S78, G91, G102, F106, and G126) and of the β1/β2 of CTDI (site II, S189 at β1 and, F201, K202, I203, G204 at β2). We also observed significant CSPs for I217 and I219 at β3, K226 at β3/β4 loop, and I231 at β4 (site I). There were also CSPs at the end of β6 (I253 and E255). It is worth noting that the triple-mutant of Sis1 (Sis1 K199N/K202N/K214N), which perturbs the α-syn binding region, does not bind the EEVD-motif^32^.

Many of the residues of CTDI with significant CSP are hydrophobic and stabilizers of the β-sandwich folding of CTDI, which is formed by the β-hairpin β2/β3 and the β-sheet (β1, β4, β5, and β6). There are two hydrophobic pockets at the β- sandwich (client-binding pocket I and II): one at the side of the pocket between β1 and β2 (pocket II) coincides with EEVD binding site II, which is the most likely client-binding site, as reported before^8,33^, and another at the opposite side, where the hydrophobic pocket is formed between β3 and β4 (pocket I), and that is likely a α-syn binding site. Note that β4 is the EEVD binding site I and that the α-syn binding site formed by β1/β2 (pocket II) showed the most pronounced CSP, therefore having the highest affinity for α-syn.

To test if EEVD competes with α-syn, we added a 10 x excess of EEVD to the α- syn/Sis1_1-352_ sample. The competition was confirmed by analysis of the residues that showed significant CSP with α-syn and further increased the CSP upon the addition of EEVD peptide (Fig 6a). Remarkably, we observed competition for residue F106 at the GF region and for residues S189, F201, K202, I203, and G204, located at the hydrophobic pocket at site II of CTDI, and residues I217, I219, and I231 located at the hydrophobic pocket at site I of CTDI. We observed competition also for residues I253 and E255 (Fig. 6a, b).

**Fig. 6.**
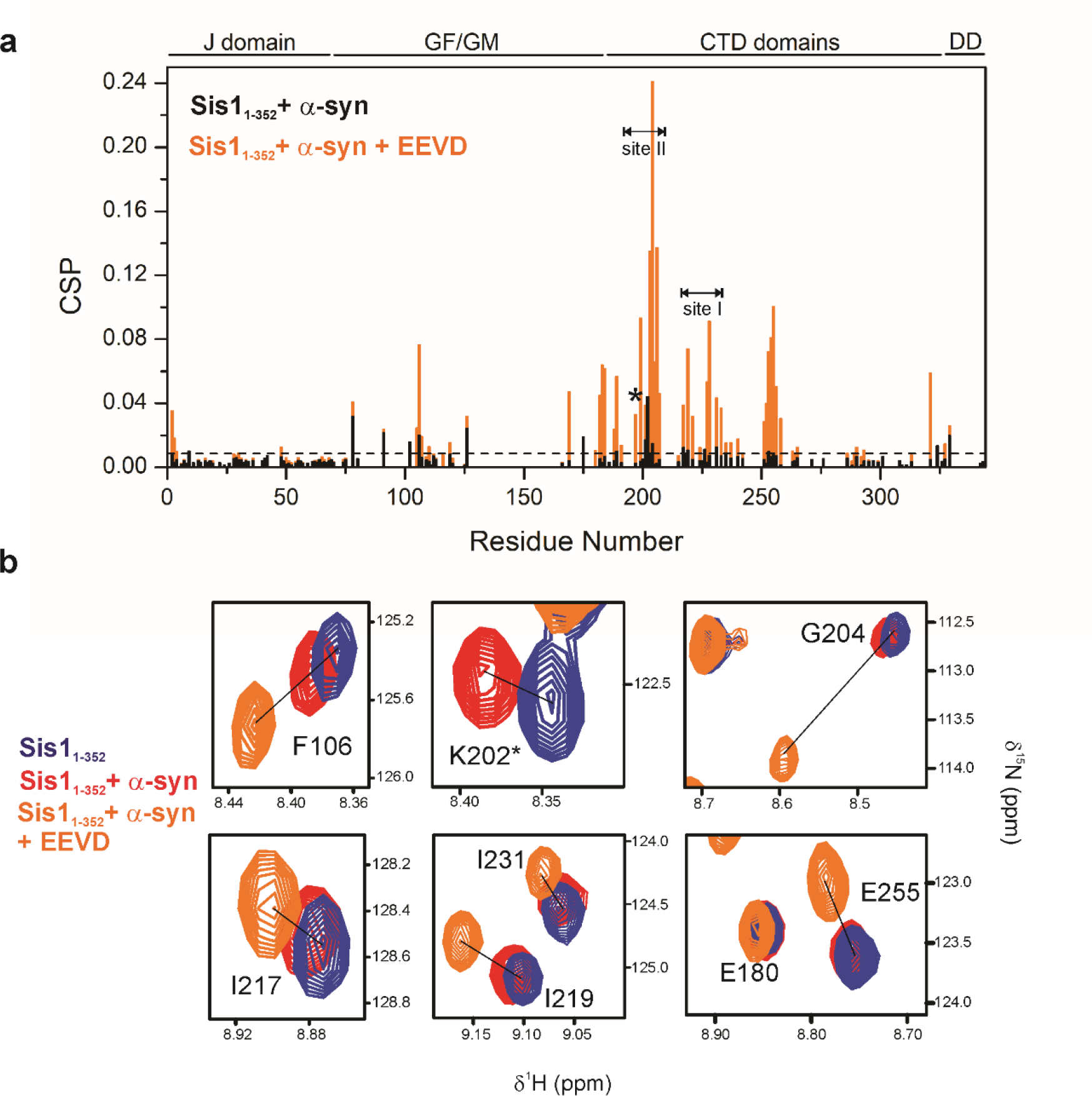
Client protein and EEVD peptide compete for the same sites of Sis1_1-352_. **a** Chemical shift perturbation (CSP) analysis of Sis1_1-352_ with α-synuclein (α-syn, black) and Sis1_1-352_ with α- syn + EEVD peptide (orange). **b** ^1^H-^15^N TROSY HSQC zoom spectra of free Sis1_1-352_ (blue) showed CSP upon interaction with α-syn (red) and α-syn + EEVD (orange) Residues that compete with the same site are shown. Sis1_1-352_ can interact simultaneously with both the client α-syn and the HSP70-EEVD peptide. *, Resonance for residue K202 disappeared upon the addition of EEVD peptide, probably due to line broadening caused by a conformational exchange.

### EEVD interaction with the GF region and the J-domain of Sis1

As discussed, the EEVD-peptide interacts transiently with the Sis1-CTDI in two sites and the analysis of both CSP and PRE data indicated interactions with residues of the J-domain (10, 14, and 42; see Fig. 3) and the GF region, mainly residues 78 to 106, which are N-terminal to α-helix 6 (Fig. 3). These data suggested that both the EEVD and the α-helix 6 competitively interact with the J-domain. The EEVD-peptide interaction with the J-domain is addressed by describing the structure of the complex between this motif and the isolated J-domain of Sis1. For that, we used the Sis1_1-81_ construct which contains the J-domain (1-72) and part of GF (73-81) and had both structure and interaction with HSP70 determined^17^. We first titrated Sis1_1-81_ with the EEVD-peptide by running a series of ^1^H-^15^N HSQCs (Fig. S11). The ^1^H-^15^N HSQC spectrum of the Sis1_1-81_:EEVD-bound is typical of a well-folded and monomeric protein and the assignment was obtained using a standard combination of triple resonance experiments. To calculate the structure of Sis1_1-81_:EEVD-bound we used ^15^N and ^13^C-edited NOESY experiments, which yielded 2750 intramolecular NOEs, enough to fold the Sis1_1-81_ (for statistics, see Table S1). We also acquired ^13^C-edited half-filtered experiments in the sample containing ^12^C-EEVD-peptide and ^13^C-Sis1_1-81_ to obtain unambiguous information on the intermolecular NOEs, but due to the transient low-affinity character of the interaction, we did not get enough intermolecular NOEs to calculate an NOE-based structure of the complex.

Upon structural calculation with Aria2.3/CNS1.21, we obtained a well-defined ensemble of 20 structures of Sis1_1-81_:EEVD-bound (Fig. S12a). The bound conformation of Sis1_1-81_ displays five α-helices (α1 to α5, Fig. S12b), four located in the J-domain (α1 to α4), and one at the beginning of the GF region (α5). The global folded conformations of α1 (residues 6-11), α2 (19-33), α3 (42-56), α4 (58-66), and α5 (69-74) are similar to those found in free Sis1_1-81_ (see reference^17^) with α2 and α3 antiparallel to one another and linked by a loop containing the HPD motif.

The ^1^H-^15^N CSP of Sis1_1-81_ upon addition of the EEVD-peptide (1:4 molar ratio) is shown in Fig. 7a. Residues with significant chemical shift differences between the free and the bound conformation are therefore identified as being directly or indirectly related to the binding site. In such a way that several Sis1_1-81_ residues had large chemical shift variations in ^1^H-^15^N-HSQC NMR spectra upon EEVD-peptide binding (Fig. 7a), delimiting a well-defined region located at the α2/α3 open patch located at the J-domain, having the interaction in the fast exchange regime on the NMR time scale (Fig. S11). In the bound state, the binding patch α2/α3 is open, when compared to the free state. α3 became slightly twisted in the bound conformation, probably to accommodate the EEVD-peptide (Fig. 7b). The RMSD between the free and bound Sis1_1-81_ states was 3.22 Å, as measured on Cα atoms. Residues A30, K32, and Y33 located at the α2, H34 and G40 located at the loop between α2 and α3, and E43, K44, F45, and I48, which are located at the α3 exhibited the largest CSP values (Fig. 7c), indicating that these residues are either engaged in direct contacts or are indirectly affected by conformational changes induced by the binding of the EEVD-peptide (Fig. 7g).

**Fig. 7.**
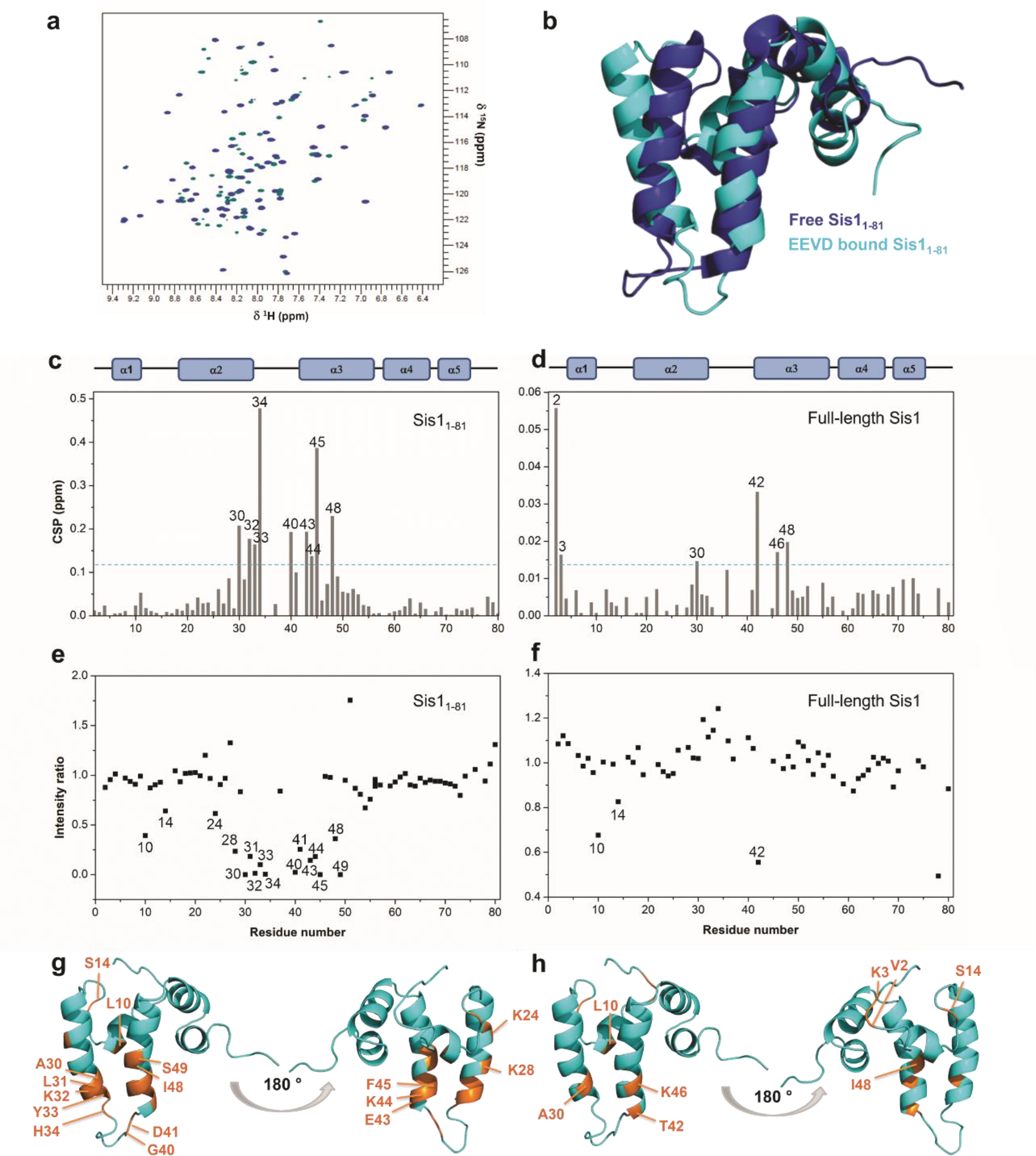
NMR analysis of Sis1_1-81_ bound to the EEVD-peptide. **a** Overlay of 2D ^1^H-^15^N HSQC spectra collected for free (blue) and EEVD-bound (4-fold molar excess; cyan) Sis1_1-81_. **b** Superposition of the NMR structures of free Sis1_1-81_ (blue, PDB 6D6X) and Sis1_1-81_:EEVD-bound (cyan, this work). **c** Chemical shift perturbation of Sis1_1-81_ resonances upon the addition of the EEVD-peptide. The CSP index is shown as vertical bars for each residue. Significant changes were considered as those with CSP larger than the average plus one standard deviation (blue dotted line, in c and d, and highlighted in orange, in g). The average and standard deviation were calculated only for the 1-81 region, different from Fig. 3 which considered the 1-352. **d** CSPs of Sis1_1-81_ resonances of the full-length Sis1 (Sis1_1-352_) upon addition of the EEVD-peptide are those above the blue line (also shown in orange in h). **e** Experimental intensity ratios of backbone amide Sis1_1-81_ in complex with the paramagnetic EEVD-peptide. **f** Experimental intensity ratios of backbone amide of the full-length Sis1 (Sis1_1-352_) in complex with the paramagnetic EEVD-peptide, however only residues 1-81 are shown to facilitate comparison with e). An intensity ratio of 1 indicates no effect of the spin label on an amide proton. CSP and PRE are mapped (orange) on the Sis1_1-81_ **g** and the Sis1_1-352_ **h** EEVD-bound conformations. In h, only residues 1-81 are shown. Note that the residues A30, at α-helix 2, and T42, K46, and I48, at α-helix 3, display either PRE or CSP for the Sis1_1-352_ suggesting the involvement of the J domain in the interaction with EEVD. Residues L10 and S14 displayed PRE for both Sis1_1-81_ and Sis1_1-352_ suggesting similar behavior for both the full-length and the isolated J domain.

The binding of EEVD to Sis1_1-81_ is much more evident and perturbs many more residues than when binding to the full-length Sis1 (Fig. 7). Residues V2 and K3 located at the N-terminal region, A30 located at the α2, and T42, K46, and I48 located at the α3 exhibited the largest CSP values (Fig. 7d). We also compared the CSPs of Sis1_1-81_ with the EEVD-bound with those of Sis1_1-81_ upon addition of HSP70, previously reported by our group^17^. Worth noting, residues A29 located at α2, H34 and T39 located at the loop, F45 and F52 located at α3, and Y67 located at α4 were perturbated by the presence of either HSP70^17^ or the EEVD-peptide, indicating that the results with peptide are similar to those with the full-length HSP70.

To get the distance restraints from the EEVD-peptide to the protein, we measured intermolecular PRE rates for the backbone amide groups of Sis1_1-81_ and Sis1_1-352_ (Fig. 3) in complex with the spin-labeled EEVD (TEMPO-EEVD, see Material and Methods). Curiously, the region affected by PRE for isolated Sis1_1-81_ was the same as observed in the CSP analysis (Fig. 7e). PRE data also showed that EEVD-peptide bound to Sis1_1-81_ is much more evident and involved many more residues than when bound to the full-length Sis1_1-352_ (Fig. 7f). Both isolated Sis1_1-81_ and the full-length Sis1_1-352_ had CSP and PRE values in the same region between α2 and α3, even though results for the full-length protein are less pronounced, probably because in the isolated domain (Sis1_1-81_) the EEVD binding site is exposed, while in the full-length Sis1_1-352_, EEVD has to compete with α- helix 6, which occludes the patch containing the EEVD binding site. The mapped interaction site with HSP70 is partially blocked by α-helix 6 in the context of the full-length Sis1, as supported by the observation of the transient interaction of the EEVD-peptide, mapped by PRE, with many residues in the GF region (78-106). Worth pointing out again, the decrease in ^15^N R_2_/R_1_ for the residues S78, G102, A105, F106, and S107 suggests a conformational selection upon EEVD-peptide binding, a condition in which the on-off equilibrium of α-helix 6 was suppressed (Fig. 2c, blue arrows). Altogether, these observations support the hypothesis that the EEVD motif competes with the α-helix 6 for the J-domain.

PRE and CSP data and HADDOCK^25,34^ were used to produce NMR data-based structural models of Sis1_1-81_:EEVD-peptide complex (see Fig. S13 for the lowest energy structural model). Ten clusters were presented for docking and the best-docked cluster was selected based on a low Haddock score and a low RMSD value (Table S2). In the Sis1_1-81_:EEVD-peptide complex model, the peptide extends between α2 and α3 with its N-terminus at the HPD-loop (Fig. S13), in good agreement with both CSP and PRE data. Molecular dynamics (MD) simulations were used to generate the Sis1_1-81_:EEVD-peptide complex and the RMSD values of the backbone atoms of the Sis1_1-81_ and EEVD-peptide from the starting structure (HADDOCK model, Fig. S13). The RMSD of the Sis1_1-81_ was stable all over the 1 μs simulation, while the RMSD of the EEVD peptide increased subtly in the first 20 ns and became stable throughout the MD simulation (Fig. 8a). The average RMSD was 4.0 ± 0.1 Å and 8.1 ± 0.2 Å for Sis1_1-81_ and EEVD-peptide, respectively (Fig. 8b). The number of contacts (distance < 0.6 nm) formed between atoms of Sis1_1-81_ and the EEVD-peptide revealed that they interacted throughout all the MD simulations with 2027 ± 339 contacts (Fig. 8b, top).

**Fig. 8.**
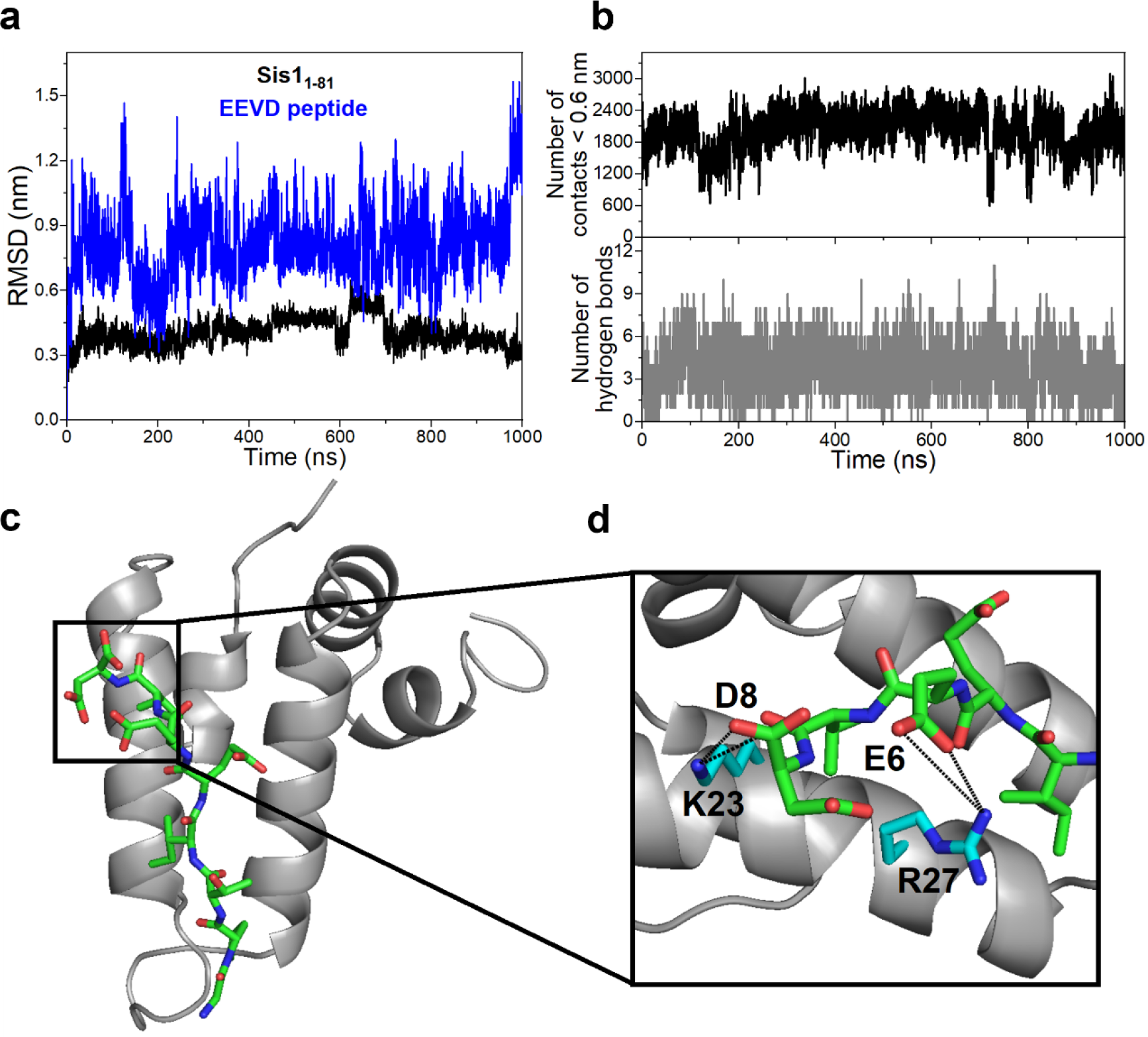
Analysis of the parameters from 1.0 μs molecular dynamics simulations of the complex between Sis1_1-81_ and EEVD peptide. **a** Values of RMSDs of the backbone atoms of Sis1_1-81_ (black) and EEVD (blue) from the starting structure (docking structure, Fig. S13). **b** number of contacts < 0.6 nm formed between atoms (Top) and a number of hydrogen bonds (Bottom) of Sis1_1-81_ and EEVD peptide. **c** Representative structure of the Sis1_1-81_:EEVD complex obtained from cluster analysis of the trajectory. Sis1_1:81_ is denoted as a gray cartoon and EEVD as sticks with carbon in green, oxygen in red, and nitrogen in blue. **d** Detail of the intermolecular salt bridges (K23-D8 and R27-E6) formed between the Sis1_1-81_ and the EEVD peptide.

The structural model showed that Sis1_1-81_:EEVD-peptide complex is formed with an average value of 4 (3.7 ± 1.5) intermolecular hydrogen bonds present throughout all the MD simulations (Fig. 8b, bottom). Percentage of persistence higher than 10% were considered and are among the residues Q20-V7, K23-D8, R27-E6, H34-P2, and E26-Y26 (Table S3). The structural model also showed that the conformation of the EEVD peptide, which is extended between α2 and α3, is in very good agreement with NMR and HADDOCK data (Fig. 8c). In addition, two salt bridges were observed between K23-D8 and R27-E6, respectively (Fig. 8d, Table S3). The formation of these specific salt bridges seems to be relevant because residues K23 and R27 are 4 residues apart from each other (i, i+4) and the same arrangement is conserved in the other two DNAJB models: DNAJB1 and DNAJB6, suggesting that they likely interact with the EEVD by the same specific salt bridges. Additionally, i, i+4 charged residues arrangement is present in some tetratricopeptide (TPR) domains stabilizing positions D(0) (D8) and E(-2) (E6) in the EEVD motifs (TPRs of Tom71, SGT2, and FKBP_92-380_, for instance). Since the TPR motif is a helix-loop-helix, as J-domain α2, loop, α3, and both structural elements bind the EEVD motif, it is conceivable they interact via similar arrangements (see for instance discussion in^35^).

## Discussion

J-domain proteins (JDPs) are responsible for transferring partially unfolded client proteins (also known as substrates) to HSP70, but the mechanism involved in this transfer is complex and still under development. This mechanism involves a series of low affinity specific transient interactions in multiple sites of both JDP (as shown here) and HSP70. NMR is a powerful method to study transient events but to fully understand the effect of multiple interactions it is necessary to work with the full-length JDP, as we accomplished here. The unique dynamical feature of Sis1 combined with the interaction of EEVD peptide and the client protein at multiple sites created the environment required for the transfer of the client to HSP70. We suggest models taking into consideration the competing and simultaneous binding of the client protein and EEVD to the globular domain CTDI, the intrinsically disordered region GF/GM, and the J domain, which may generate entropic forces able to orient the transfer of the client to HSP70, a mechanism observed for other proteins^36^.

Fig. 9a illustrates a minimalist, as it captures the intrinsic nature of interactions, as well as the most accepted mechanism for the JDPs/HSP70 cycle^5,31,37^. First, JDP recognizes and binds to the unfolded client protein. The next step is the binding of the binary complex JDP/client to the HSP70. The ternary complex client/JDP/HSP70(ATP) provides the necessary environment for the transfer of the client unfolded protein from the JDP to the HSP70. JDP interacts with two sites of HSP70: (i) the intrinsically disordered C-terminal that contains the EEVD motif (the focus of this work), and (ii) the interface between the nucleotide-binding domain (NBD) and the substrate binding domain (SBD). The interaction of the J-domain of the JDP with the NBD/SBD interface stimulates the ATPase activity leading to the dissociation of the JDP and the formation of a binary complex client/HSP70(ADP). At this step, the HSP70 structure changes from the open to the closed conformational state, creating the microenvironment for client refolding. The cycle is then completed with the binding of the nucleotide exchanging factor (NEF), which promotes the release of the ADP, folded client protein, and further dissociation of the NEF itself. At this stage, the HSP70(ATP) is regenerated, being apt to bind to another JDP carrying an unfolded client protein.

**Fig. 9.**
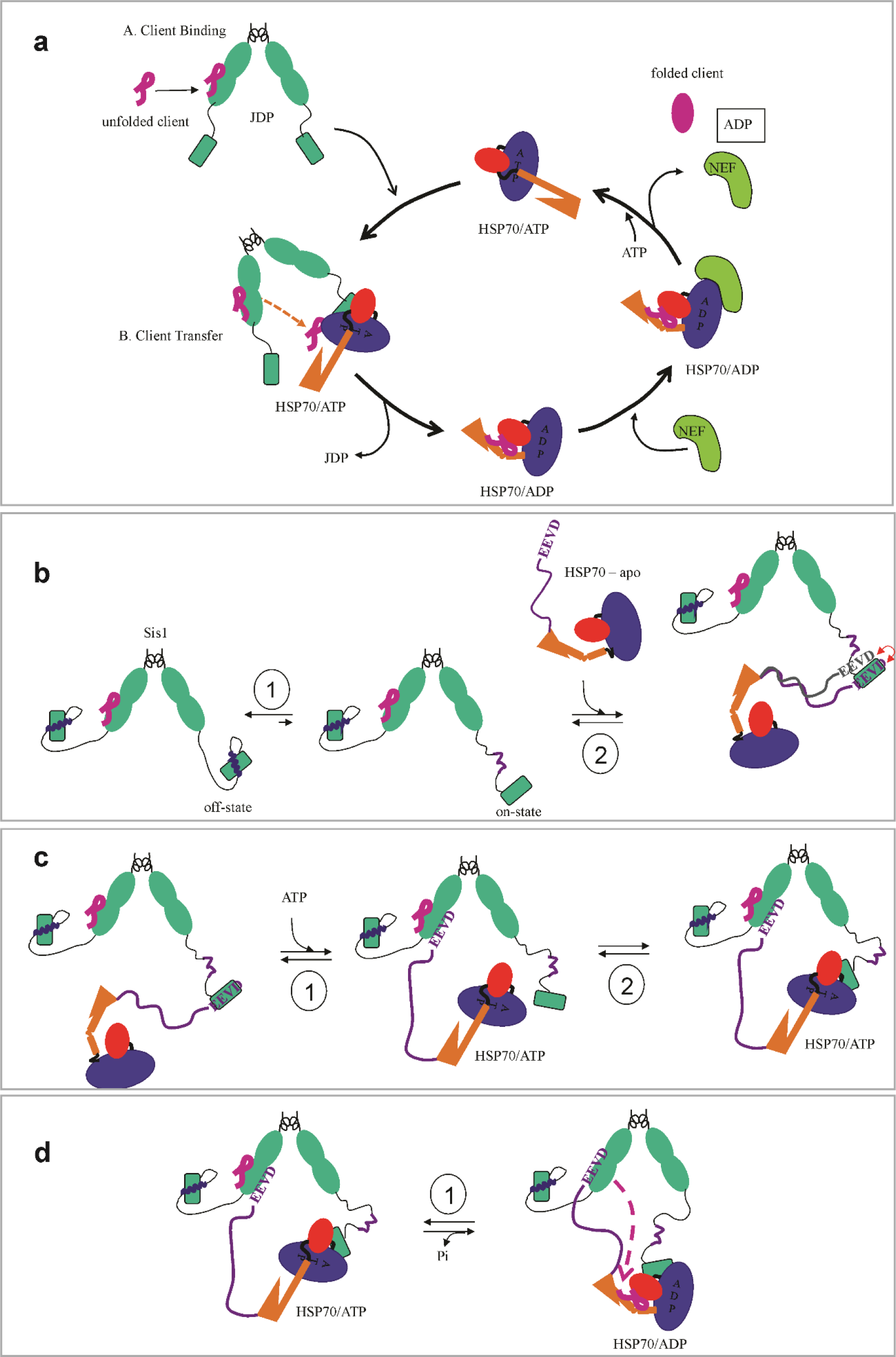
Chaperone cycle of HSP70 and class B JDP. Fig. S14 shows that the schematic drawings of HSP70 here presented are realistic, following the crystallographic structures of conformational states of HSP70. **a** Schematic drawing of the most accepted mechanism for the JDPs/HSP70 cycle^5,31,37^. JDPs (green) recognize and binds to a nascent or misfolded protein (client protein, magenta) through their peptide-binding domain and interact with an ATP-bound Hsp70 through their J domain. Association of the J domain at the interface between the nucleotide-binding domain (NBD) and the substrate binding domain (SBD) stimulates the ATP hydrolysis, promoting a conformational change in HSP70 and transition to the ADP-bound state, which has a high affinity for the substrate. JDP then leaves the complex and the nucleotide exchange factor (NEF) induces ADP dissociation. If the client protein achieves the native state (folded client), the cycle ends; if it remains unfolded, the JDP rebinds to it and the cycle begins again; **b** Step 1 describes the on-off equilibrium between the J-associated (off-state) and J-dissociated (on-state) α-helix 6. The on-state exposes the recognition patch at the J-domain, susceptible to association with HSP70-EEVD. Step 2 illustrates the transient binding of the C-terminal EEVD of the apo state of HSP70 with the GF region and J-domain. Note that the client protein preferentially binds to pocket II (site II); **c** Step 1 illustrates the EEVD binding to site I of CTDI, while the client is bound to pocket II (site II). ATP binds to NBD causing HSP70 conformational change towards the open state. In this step, the J-domain becomes available to bind to the open conformation of HSP70/ATP. Step 2 shows the binding of the J-domain at the interface between NBD (blue) and SBDβ (red). **d** Next SBD interacts with ultra-affinity to the client protein, with the concomitant aid of the competitive action of EEVD that dissociates from the client protein and binds to the site II of CTDI. The purple arrow illustrates the transfer of the client protein from CTDI to the closed SBD/ADP since the ATPase activity of the NBD is activated to its maximum by the combined action of the J-domain association and the binding of the client protein.

In this work, we carried out solution studies of the interaction of the EEVD-peptide with the full-length Sis1 and showed that EEVD binds in multiple sites, at the J, GF, and CTDI domains. We also showed that the client α-syn elicited significant CSP at three sites, one at the GF/GM intrinsically disordered region and two at CTD1 (pockets I and II). Additionally, of high relevance, our results strongly suggest that EEVD competes with the client protein for the GF and CTDI domains, preferentially for site II (client binding pocket II). Several studies of the interaction of EEVD with JDPs, from other research groups, were important to show the interaction at the CTD domain^13,27,38^ (CTDI for Sis1), although working with the truncated form of the protein (CTDI/CTDII only). A major challenge in structural biology is the determination of transient events and this work corroborates the complexity of the events involving the interaction of Sis1 with the HSP70 C-terminal in multiple sites. All described interactions with either the EEVD motif or the client protein were transient and specific, which are intrinsic to low-affinity binding. They should not be mistaken for non-specific interaction which would not elicit combined CSP and PRE effects. To generate a more complete understanding of this interaction, this work provided a thorough investigation of the structure and the role of the dynamics of the full-length Sis1_1-352_, and on the Sis1_1-81_ construct, bound to an EEVD-peptide.

We addressed the effect of these interactions on the protein dynamics by measuring relaxation parameters and pin-pointing the EEVD-peptide interacting sites using CSP and PRE data combined with an experimental-driven computational docking. The EEVD-peptide was able to bind transiently to multiple sites in the full-length Sis1, two of them at the CTDI, at sites I and II, one at the boundary of α-helix 6, in the GF-region and the other at the J-domain. Interaction of EEVD with site I was previously observed in the crystal structure of the complex with an EEVD-peptide with DNAJB1^27^. For Sis1, this interaction was first reported here. The interaction of the EEVD with the site II of Sis1-CTDI was observed previously in the crystal structure of the complex Sis1:EEVD (actually a construct containing only the CTDI/CTDII domains)^13^, and in solution by CSP also using a construct of only the CTDI/CTDII domains^38^. Different from what occurs with Sis1, the EEVD binds at the CTDII of a *T. thermophilus* JDP (ttHsp40)_38._

Next, we will discuss and propose models (Fig. 9b-d) that illustrate how the labile interaction at multiple sites can contribute to the understanding of the mechanism of the client transfer from Sis1 to HSP70. These models are supported by interaction studies and the changes in Sis1 dynamics, and they should be taken into consideration in further studies. They send the message that labile interaction in multiple sites plays key roles, especially when they involve intrinsically disordered regions where entropic forces are involved^36^.

A new information for the mechanistic path of Sis1/HSP70 interaction is the observed interaction of the EEVD-peptide with the J-domain and the GF region. Fig. 9b illustrates how the interaction at these regions could contribute to the regulation of the JDP/HSP70 cycle. This work presents robust experimental evidence on the binding of EEVD-peptide with the Sis1-GF domain (CSP and PRE, Fig. 3). Nevertheless, only a few residues at the J-domain in the full-length Sis1 had CSP and PRE effect. On the other hand, for the isolated Sis1-J-domain, we provided richer and more detailed structural information on the complex with EEVD-peptide (Fig. 7 and 8). We interpreted the difference between the EEVD binding to the isolated J domain and in the context of the full-length protein, taking into consideration the reported autoinhibition mechanism described for human DNAJB1_1-111_, in which the J-domain is complexed with the α-helix 5^15^ (α-helix 6 for Sis1). Alphafold^21^ prediction of the structure of Sis1 (Fig. 5) suggested the association of α-helix 6 with the J domain, hiding the EEVD interaction patch. We suggest that the interaction of the EEVD-peptide with Sis1 is modulated by an on-off equilibrium between the J domain-dissociated (on-state) and J domain-associated (off-state) α-helix 6, in an autoinhibition mechanism. The on-off equilibrium is illustrated in path 1 of Fig. 9b. The on-state exposes the EEVD interaction patch of the J-domain that is composed of α-helix 2 and 3 and the loop between them, which contains the HPD motif. Remarkably, the same J-domain recognition patch binds to the EEVD motif, α- helix 6^15^, and to the interface between the NBD/SBDβ of HSP70^17,28^. The suggested mechanism described in Fig. 9b contemplates a labile anchoring of HSP70, through the EEVD motif to the J and GF domains of Sis1 (red double arrow in Fig. 9b). The intrinsically disordered C-terminal of HSP70 would aid in extending the range of interaction to JDP, approximating the HSP70 to Sis1 and favoring the further binding of the J-domain to the NBD/SBDβ interface. It could also favor the on-off equilibrium toward an on-state (Fig. 9b step 1), in which the J domain binding patch is ready to interact with HSP70, stimulating the ATPase activity (Fig. 9c).

Supporting the suggested model are the following data. The J-domain interacting patch is exposed in the isolated J domain (Sis_1-81_), while it is mostly occluded in the full-length Sis_1-352_. The PRE and CSP data for this patch was much more pronounced when analyzing its binding to Sis_1-81_ than to Sis_1-352_ (Fig. 9), although for the Sis_1-352_, CSP, and PRE were present for the J-domain recognition patch: A30 at α-helix 2, T42, K46, and I48, at α-helix 3. Residues L10 and S14 displayed PRE for both Sis1_1-81_ and Sis1_1-352_ suggesting similar behavior for both the full-length and the isolated J domain (Fig 7). Additionally, the CSP and PRE were much more robust for the adjacencies of α-helix 6, located at the GF domain, when Sis_1-352_ was bound to the EEVD-peptide. We interpreted these data as 1) the interaction of EEVD with GF can occur with both the on-and off-states; 2) only when α-helix 6 is dissociated from the J-domain (on-state) the EEVD binds to the interaction patch of the J-domain (Fig. 9b step 2). These interpretations are supported by the suppression of conformational exchange in the residues of GF region upon binding to the EEVD peptide. Furthermore, there was a subtle increase in the rotational freedom of the J-domain in the presence of EEVD. The higher rotational freedom of the J-domain contributes to the biological mechanism of the JDPs, and the binding of EEVD makes it more available for interactions. Further studies are necessary to understand the interplay between the EEVD binding to the J-domain interacting patch and the GF region.

The EEVD/J-domain association is mostly electrostatic, with two salt bridges. On the other hand, the GF region (78-105) has no charged residues, and EEVD probably interacts with GF (presence of PRE, Fig. 3) via hydrophobic contacts involving its residues G1, P2, T3, I4, and V7. Since the α-helix 6 association with the J-domain binding patch is also mediated by hydrophobic interaction, through its residues A111, I114, F115, F118, and F119, we suggest that the EEVD association with GF favors EEVD to compete with a-helix 6 to the J-domain interacting patch. To sum up, the competing interaction of EEVD with the J-domain patch shifts the equilibrium to the on-state (Fig. 9b, step 2, red double arrow), exposing the J-domain binding site to interact with the NBD and SBDβ of the open/closed conformation of HSP70.

We described the interaction of the EEVD-peptide with both sites I and II of Sis1-CTDI. Site I is described as more specific toward the binding site of the EEVD motif^27^ while binding with lower affinity to α-syn (CSP is larger for site II, Fig. 6), from which it raises competitivity for this site. In this model, EEVD peptide competes preferentially for the client binding site (pocket II, Fig. S15), which is the major hydrophobic site for the binding of the client protein^8,38^. There is no mechanism describing whether each protomer of JDP may or may not simultaneously access different binders. For instance, Suzuki and Cols (2010)^27^ suggested for DNAJB1 that the mechanism involves the two protomers of the JDP. The client would bind to one protomer at site II, while the EEVD motif would bind to the other but at site I. Our data mapped multiple points of EEVD interaction, which allow some predictions. Fig 9c presents a model that contemplates the two vicinal EEVD binding sites with the client and EEVD to the same protomer. In this model, step 1 (Fig. 9c), predicts that the two protomers are recruited by the HSP70 conformational change triggered by ATP. The EEVD associated with the J domain in one protomer could be transferred to the CTDI in the other protomer and associated at site I. Thus, the conformational change in HSP70 inverts the position of the C-terminal tail that contains the EEVD motif. Step 1 prepares for step 2 (Fig. 9c), which creates a conformational state in which HSP70 is anchored by two sites. One to the J domain, interacting with the NBD/SBDβ interface, which is also triggered by ATP and J domain, and the other at site I of CTDI using the EEVD motif of the C-terminal tail. We would like to point out that this model is consistent with the multiple interaction points, however, a different model may also explain some of the results.

The competition at multiple sites probably plays important role in the still-unknown transfer mechanism of the client from CTDI pocket II of JDP to the SBD of HSP70 since the EEVD peptide competes for all client interaction sites, CTDI pockets I and II, and GF/GM intrinsically disordered region. We suggest that the conformational state in that HSP70 is anchored in two sites of JDP enables the creation of a favorable environment for client transfer (Fig 9d). As observed here, the client protein interacts with CTDI, but also with the GF/GM region. We should keep in mind that client proteins, such as α-syn, are relatively large unfolded or partially unfolded polypeptides that could make several simultaneous interactions with the JDP. The simultaneous binding of the client protein to the globular domain CTDI and the intrinsically disordered region GF/GM may generate entropic forces to orient the transfer, as demonstrated to other proteins^36^.

Finally, all interactions of the EEVD with sites I and II of the CTDI are of low affinity and largely electrostatic (salt bridges and hydrogen bonds) raising the hypothesis that the EEVD has two functions: (i) binding to site I of CTDI, keeping HSP70 anchored to Sis1 even without the J-domain association with NBD/SBDβ (Fig 9c) and (ii) competing with the client protein for pocket II (EEVD binding site II) (Fig. 6 and 9d), helping in the dissociation of the client protein, which is transferred to the ultra-affinity state of HSP70. The ternary complex JDP:HSP70:client dissociates from the JDP, consequently releasing HSP70:client complex. Altogether, the EEVD interacts with multiple sites of Sis1_1-352_ acting as a modulating agent or a factor of disorder, facilitating the exposure and availability of the J-domain and CTDI to the interaction with HSP70.

## Methods

### Sample Preparation

Sis1_1-352_ (UNIPROT P25294) and Sis1_1-81_ expression and purification were carried out according to previous work^39,40^. GPTIEEVD and CGPTIEEVD peptides referring to the C-terminal motif of HSP70 were synthesized and purified by GenOne Biotechnologies (Rio de Janeiro, Brazil).

### NMR spectroscopy

The NMR experiments to assign Sis1_1-352_ were collected in a Bruker Avance III HD 950 MHz Spectrometer equipped with ^13^C, ^15^N, ^1^H, ^2^H TXI cryoprobe. Sis1_1-352_:EEVD-bound, Sis1_1-81_ and Sis1_1-81_:EEVD-bound were collected on a Bruker Avance III HD 900 MHz spectrometer equipped with an inverse-detection triple resonance z-gradient TXI probe. All NMR samples contained 300 μM of Sis1_1-352_ or 1 mM of Sis1_1-81_ in 25 mM Tris-HCl pH 7.5, 200 mM NaCl and 10 % D_2_O at 303K or 298 K, respectively. NMR data were processed and analyzed with nmrPipe^41^ and CcpNmr Analysis^42^ available on the NMRbox platform^43^.

### Backbone dynamics

^15^N backbone amide relaxation parameters (R_1_ and R_2_) were acquired on Bruker 900 MHz for a ^15^N-Sis1_1-352_ at 303 K. The TROSY-based ^15^N longitudinal (R_1_) relaxation rates were measured at 20, 300, and 600 ms, and transverse (R_2_) relaxation rates were measured at 8.65, 16.96, and 33.92 ms. Relaxation parameters were determined by fitting T_1_ and T_2_ peak intensities to a single exponential decay. ^15^N R_1_ and R_2_ values were determined from the fit of the peak intensities using a mono-exponential equation. Experimental errors of the relaxation parameters were evaluated based on the signal/noise ratio as described previously^44^. All relaxation experiments were acquired as pseudo-3D spectra and converted to 2D data sets. NMR spectra were processed and analyzed with NMRPipe^41^. The overall rotational correlation time (τ_c_) of Sis1_1-81_ was estimated from the mean values of R_1_ and R_2_ measured for each domain, as follows: 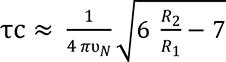, where ʋ_N_ is the ^15^N resonance frequency (Hz), R_1_ and R_2_ are the mean values of the ^15^N relaxation rates^45^.

### Chemical shift assignment

The backbone resonance assignments of Sis1_1-352_, data were acquired from the following two-dimensional (2D) and three-dimensional (3D) NMR experiments: all BEST-TROSY versions^46^: BT-HSQC, BT-^13^C-HSQC, BT-HNCACB, BT-HNCA, BT-HNCACO, BT-HNCO, BT-HNCOCA, BT-HNCOCACB, BT-N-NOESY, and BT-HNCANH. The assigned chemical shift values of backbone ^15^N, ^13^Cα, and ^13^C’ of Sis1_1-352_ were deposited in the BMRB (ID 51817) and used as input for the TALOS-N program^19,20^ to predict secondary structures. The backbone and side chain assignments of Sis1_1-81_:EEVD-bound were also deposited in the BMRB (ID 51187).

### Site-direct Spin-Labeling experiments

A 20-fold excess of 4-maleimide-TEMPO (Sigma-Aldrich) dissolved in acetonitrile was added to CGPTIEEVD peptide at 1 mM in 25 mM Tris-HCl pH 7.5, 200 mM NaCl and incubation for 2 h at room temperature. The final product was the TEMPO spin label attached to the reactive thiol functional group of Cys1. To remove unreacted and excess maleimide spin label, the reaction mixture was injected into a C18 column and HPLC chromatography (Cytiva) in acetonitrile and water and eluted with a 30-70 % (water:acetonitrile 0.1 % TFA) of gradient

### Paramagnetic relaxation enhancement (PRE)

The direct interaction of Sis1_1-352_ and Sis1_1-81_ with EEVD motif was investigated by analyzing the PRE with the TEMPO-labeled EEVD peptide. ^15^N-Sis1_1-81_ was titrated with TEMPO-labeled CGPTIEEVD peptide (paramagnetic sample) and ^1^H-^15^N TROSY (for Sis1_1-352_) or HSQC (for Sis1_1-81_) spectra were recorded at 900 MHz. The diamagnetic samples were obtained after the reduction with 3 mM ascorbic acid, at room temperature for 4 h. PRE was obtained by the intensity ratio between the paramagnetic and diamagnetic spectrum (*PRE* = *αI^para^*/*I^dia^*). We observed PRE effects higher than one because of the larger longitudinal relaxation time for the diamagnetic sample when compared to the paramagnetic and a short relaxation time used in the TROSY or HSQC spectra (1 and 1.3 s, respectively). Because of that, we introduced a correction factor (α) to make the average <PRE> =1 for the residues distant from the paramagnetic center.

To calibrate the PRE effect into semiquantitative distance ranges we considered a linear change of the PRE effect as a function of the distance from each NH to the paramagnetic center (d) between 12 and 30 Å^24,47^. We considered d >30 Å for PRE =1 and d < 12 Å for PRE = 0. Distances between 12 and 30 Å were calculated (d_calc_) assuming a linear change proportional to PRE. The distances were used semi-quantitatively for the Haddock calculation: For PRE = 0, d < 12 Å, and for PRE between 0.8 and 0, d was in the range between d_calc_ and 30 Å.

### Chemical shift mapping

To identify the interaction of Sis1_1-352_ or Sis1_1-81_ with EEVD peptide, ^1^H-^15^N-TROSY spectra were collected for free ^15^N-labeled Sis1_1-352_ at 0.3 mM and upon addition of GPTIEEVD- (from now on referred as EEVD) (1:4 protein: peptide). For ^15^N-Sis1_1-81_ at 0.5 mM, ^1^H-^15^N HSQC spectra were collected of free and upon addition of EEVD (1:4 protein: peptide). For the interaction of Sis1_1-352_ with α-synuclein (α-syn) as a client protein, ^1^H-^15^N-TROSY spectra were collected for 0.3 mM of free ^15^N-labeled Sis1_1-352_, and after the addition of 0.3 mM of α-syn. To the same sample, a ∼10-fold excess of EEVD, related to Sis1, was added. Chemical shift perturbations (CSP) were calculated for each backbone amide group, as follows: 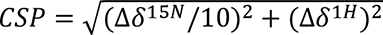, where Δδ^15N^ and Δδ^1H^ are the chemical shift differences between the free and bound states of Sis1_1-81_:EEVD or Sis1_1-352_:EEVD. CSP greater than 1 standard deviation from the mean was considered significant.

### Structure calculation of Sis1_1-81_:EEVD

NMR data for structure calculation of Sis1_1-81_:EEVD were collected on a 1 mM ^13^C/^15^N-labeled Sis1_1–81_ in 25 mM Tris-HCl pH 7.5, 200 mM NaCl and 10 % D_2_O at 900 MHz. Distance restraints were derived from 3D ^15^N-NOESY-HSQC and 3D ^13^C-NOESY-HSQC both aliphatic and aromatic and dihedral angle restraints were derived from ^1^HN, ^15^N, ^13^Cα, ^1^Hα, ^13^Cβ, and C’ chemical shifts using TALOS-N^20^. The amino acid sequence, the chemical shift lists, the dihedral angle values, and the three NOESY data sets were used as input files for structural determination.

Three-dimensional structures of the Sis1_1-81_:EEVD-bound were calculated using the program Aria 2.3^48,49^ combined with CNS 1.2^50^ available on the NMRbox platform ^43^. In the final calculation the 20 lowest-energy structures, derived from the water refinement step, were selected as representative of the ensemble of Sis1_1-81_:EEVD conformations in solution. For quality validation, we used Protein Structure Validation Software suite (PSVS)^51^ (https://montelionelab.chem.rpi.edu/PSVS/PSVS/). Structures were visualized with PyMOL^52^ and the atomic coordinates of Sis1_1-81_:EEVD were deposited in the *Protein Data Bank* under accession code 8EOD.

### Molecular Docking and Molecular Dynamic (MD) Simulations

The software Haddock^25^ was used to dock the EEVD peptide to CTDI (residues 180 to 257) of Sis1_1-352_ using the residues with CSP (Fig. 3b) as active residues and distance restraints semi-quantitatively calibrated from the PRE data (Table S3). Passive residues were automatically assigned as those surrounding the active ones. Residues with low order parameters (S^2^ < 0.6, Fig. 2B) were set as fully flexible. For Sis1_1-352_, we have NMR-based interaction information with many regions but created docking models for CTDI only. It was unfeasible to create an interaction model of one EEVD peptide that was compatible with all the experimental data. The HADDOCK (version 2.4) server (https://wenmr.science.uu.nl/haddock2.4/)^25^ was used for the modeling of CTDI:EEVD. The protein structural coordinates of CTDI used as input was obtained from an Alphafold prediction, which is almost identical to the Protein Data Bank (PDB) under access code 2B26^13^. In total, 2000 complex structures of rigid-body docking were calculated by using the standard HADDOCK protocol with an optimized potential for liquid simulation (OPLSX) parameters. The final 200 lowest-energy structures were selected for subsequent explicit solvent (water) and semi-flexible simulated annealing refinement, to optimize side chain constants. The Volmap tool of the Visual Molecular Dynamics (VMD)^26^ software was used for the construction of the atomistic probability map of CTDI-Sis1:EEVD. An ensemble of 100 structures lowest energy structures calculated from Haddock was used. The 100 structures were clustered into two groups. Cluster I with all the poses binding to Site I and Cluster II with all the poses binding to Site II (Figure S1).

The software Haddock^25^ was used to dock the EEVD peptide to the EEVD-bound conformation of Sis1_1-81_ (Fig.S12, PDB 8EOD) using the residues with CSP (Fig. 7c) as active residues and distance restraints semi-quantitatively calibrated from the PRE data (Table S5). Passive residues were automatically assigned as those surrounding the active ones. The HADDOCK (version 2.4) server (https://wenmr.science.uu.nl/haddock2.4/ was used for the modeling of Sis1_1-81_:EEVD. In total, 2000 complex structures of rigid-body docking were calculated by using the standard HADDOCK protocol with an optimized potential for liquid simulation (OPLSX) parameters. The final 200 lowest-energy structures were selected for subsequent explicit solvent (water) and semi-flexible simulated annealing refinement, to optimize side chain constants. The final structures were clustered using the fraction of common contacts (FCC) with a cutoff of 0.6.

Molecular dynamics (MD) calculations for docking models of CTDI at site I and site II and the 2 lowest energy structure of the highest representative cluster of Sis1_1-81_:EEVD complex were performed using GROMACS (version 5.1.4) ^53^. The molecular systems were modeled with the corrected AMBER14-OL15 package, including the ff14sb protein force field ^54^, as well as the TIP3P water model ^55^. The structural models of the complexes (from molecular docking) were placed in the center of a cubic box solvated by a solution of 200 mM NaCl in water. Periodic boundary conditions were used, and all simulations were performed in an NPT ensemble, keeping the system at 25 °C and 1.0 bar using Nose-Hoover thermostat (τ_T_ = 2 ps) and Parrinello-Rahman barostat (τ_P_ = 2 ps and compressibility = 4.5×10^−5^·bar^−1^). A cutoff of 12 Å for both Lennard-Jones and Coulomb potentials was used. The long-range electrostatic interactions were calculated using the particle mesh Ewald (PME) algorithm. A conjugate gradient minimization algorithm was used to relax the superposition of atoms generated in the box construction process. Energy minimizations were carried out with the steepest descent integrator and conjugate gradient algorithm, using 1,000 kJ·mol^−1^·nm^−1^ as the maximum force criterion. Five hundred thousand steps of molecular dynamics were performed for NVT and NPT equilibration, applying force constants of 1,000 kJ·mol^−1^·nm^−2^ to all heavy atoms of the complex. At the end of preparation, a 100 ns MD pulling simulation of the molecular system was carried out using a spring constant of 1,000 kJ·mol^−1^·nm^−2^ between the protein and the peptide. Next, 1 μs MD simulation was performed for data acquisition. Following the dynamic, the trajectories of the complex were firstly concatenated and analyzed according to the RMSD for the backbone atoms of protein and peptide, many contacts for distances lower than 0.6 nm between pairs of atoms of CTDI-Sis1/Sis1_1-81_ and EEVD peptide, and several protein-peptide hydrogen bonds with cutoff distance (heavy atoms) of 3.5 Å and maximum angle of 30°. The percentages of protein-peptide hydrogen bond persistence were obtained from *plot_hbmap_generic.pl* script^56^. The number of protein-peptide hydrogen bonds with persistence greater than 10% was considered. The tool *g_cluster* of GROMACS^57^ package was used to perform cluster analysis in the 1.0 μs MD trajectory of the complex using a cutoff of 3 Å. The structure of the first cluster was used as a representative structure of the complex The structural representation of the constructed model was displayed using PyMOL^52^.

## Abbreviations

CSP: chemical shift perturbation
CTD: C-Terminal Domain
HSP: heat shock protein
HSP70: 70 kDa heat shock protein
IDR: intrinsically disordered region
JDP: J-domain protein
NBD: nucleotide-binding domain
R_1_: longitudinal relaxation rate
PRE: Paramagnetic relaxation enhancement
R_2_: transverse relaxation rate
SBD: substrate binding domain
RMSD: root-mean-square deviation
Sis1: type B JDP from *Saccharomyces cerevisiae*
Ssa1: HSP70 from *Saccharomyces cerevisiae*

## Acknowledgements

The authors thank Dr. Bernhard Brutsche (responsible scientist), Dr. Adrien Favier (NMR platform engineer), Dr Jérôme Boisbouvier (Researcher) for help with the acquisition of NMR data. This work used the platforms of the Grenoble Instruct-ERIC center (ISBG; UAR 3518 CNRS-CEA-UGA-EMBL) within the Grenoble Partnership for Structural Biology (PSB), with financial support from the TGIR-RMN-THC Fr3050 CNRS and the National Center of Nuclear Magnetic Resonance (CNRMN/UFRJ, https://www.cenabio.ufrj.br/index.php). This work has been supported by iNEXT-Discovery (871037), funded by the Horizon 2020 program of the European Commission and FAPESP (2012/50161-8; 2017/26131-5). CHIR and FCLA have research fellowships from CNPq (305148-2019-2 and 313517/2021-5, respectively). FCLA have a research fellowship of FAPERJ (273303, 204432 and 267010). The following authors received a research fellowship from FAPESP: COM (2019/16114-1), and GMSP (2017/01074-9; 2018/11948-9).

## Author contributions

Data collection, analysis, and interpretation: COM, GMSP, FCLA, GCA, IPC. Data analysis and interpretation and design of the work: CHIR.; FCLA. This work results from the collaboration of two research groups: Laboratory of Biochemistry of the Chaperome (PI CHIR) and Biomolecular NMR Laboratory (PI FCLA). All authors: drafted and critically reviewed the article.

## Competing interest statement

The authors declare no competing interests.

## Notes

### Competing Interest Statement

The authors have declared no competing interest.

### Summary of Updates

Abstract updated

