## Supplementary Material for "Structural and dynamical basis for the interaction of HSP70-EEVD with JDP Sis1"

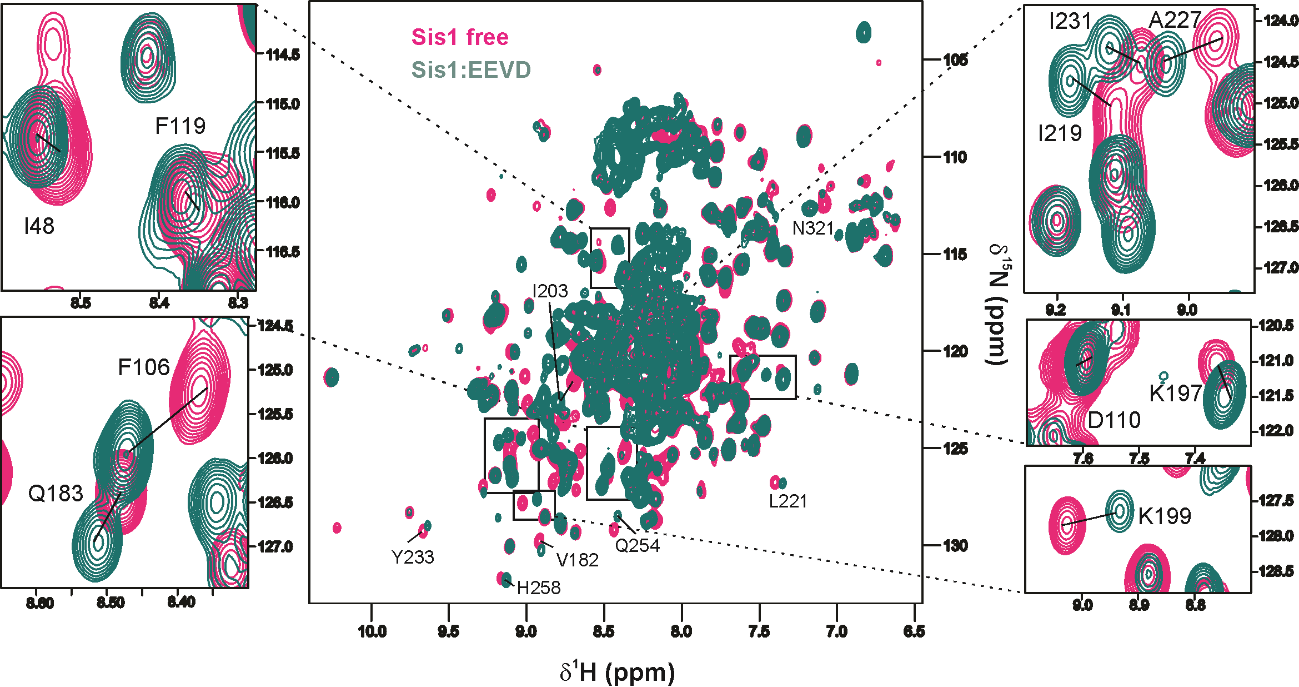

**Fig S1. |** **Overlay of ^1^H,^15^N TROSY HSQC spectra acquired for full-length Sis1 (^15^N-Sis1_1-352_) with an excess of EEVD-peptide (1:4)**. Sis1_1-352_ in the absence (magenta) and presence (green) of EEVD peptide. Full spectra are shown in the middle, highlighting residue I48 (J-domain); F106, D110, and F119 (GF region) and, Q183, K197, K199, I219, A227, and I231 (CTDI); all perturbated in the presence of EEVD indicating their participation in the interaction.

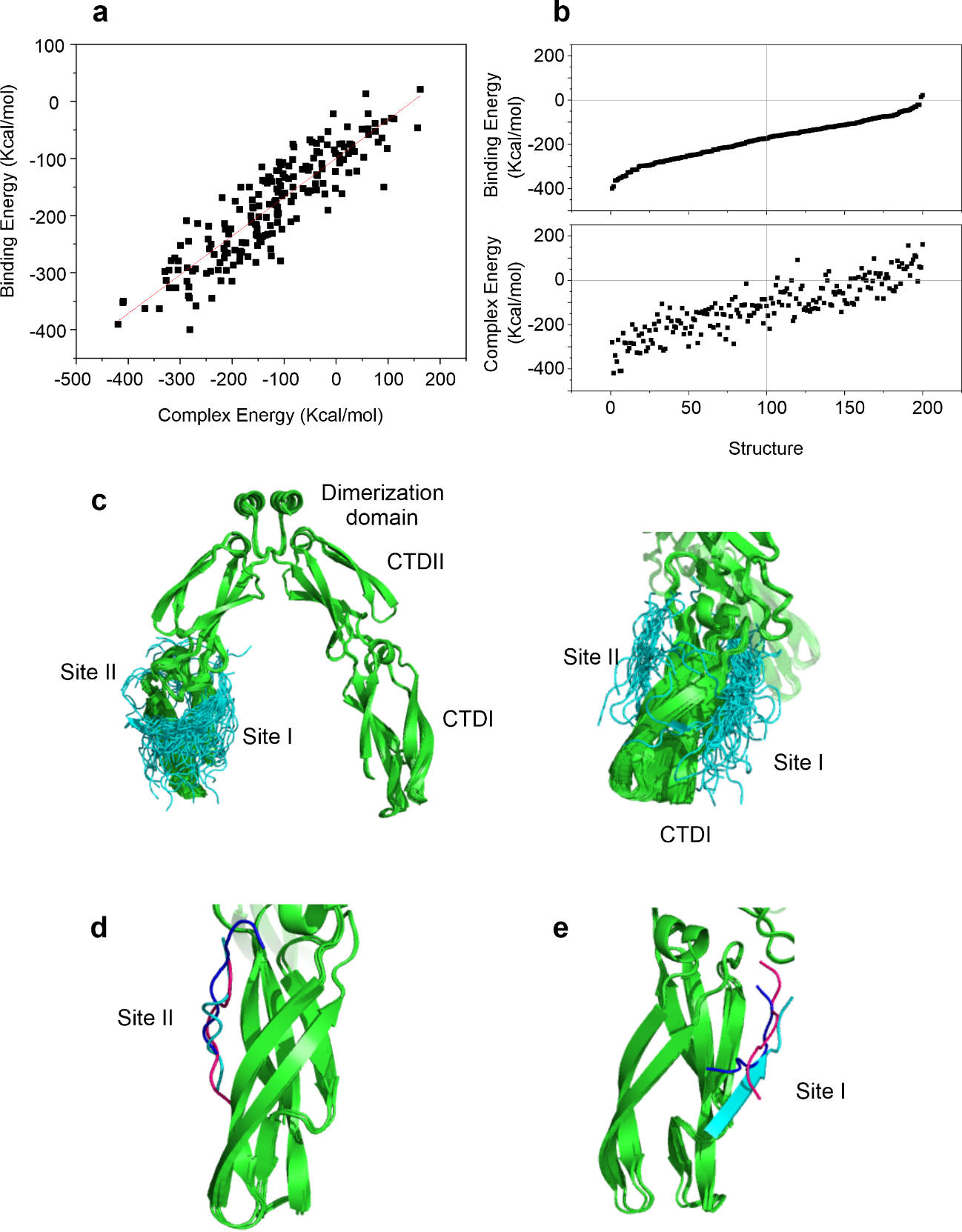

**Fig. S2 | Analysis of 200 lowest energy structures calculated from Haddock using PRE and CSP restraints. a** Binding energy as a function of complex energy. **b** Binding energy and the energy of the CTDI:EEVD complex as a function of the 200 docked structures. Note that the 100 lowest energy structures had both negative binding and CTDI:EEVD complex energy. We chose to construct the mass density plot from the clusters at site I and II of these 100 energy poses. **c** Cartoon representation of the superposition of the 100 lowest binding energy poses bound to sites I and II of CTDI. EEVD peptide is in cyan and Sis1 in green. **d** Cartoon representation of the superposition of the 3 lowest binding energy of the EEVD peptide (cyan, dark blue and pink) bound to site II and **e** Cartoon representation of the superposition of the 3 lowest binding energy of the EEVD peptide (cyan, dark blue and pink) bound to site I.

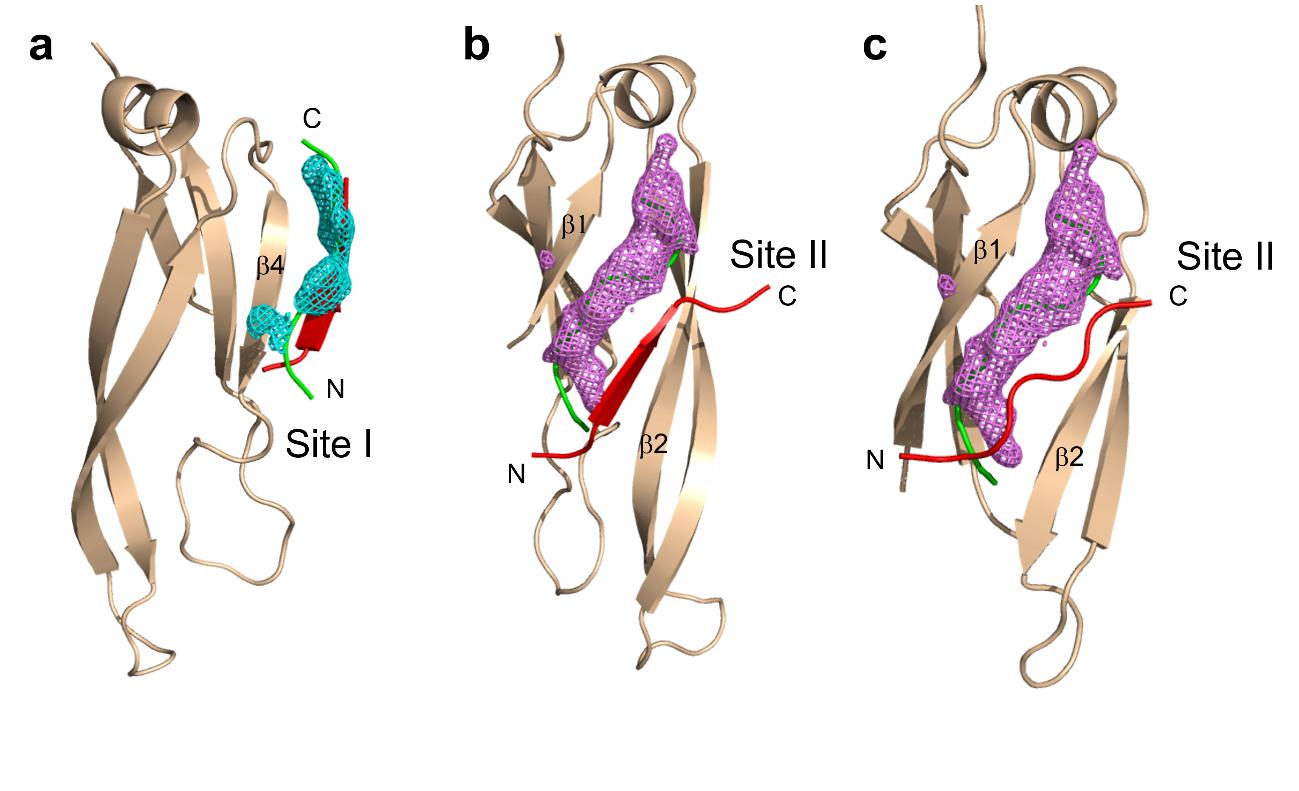
**Fig. S3 | EEVD-peptide and interaction with the binding sites in the crystal structures of DNAJB1 and Sis1.** Cartoon representation of the CTDI (wheat color) from **a** DNAJB1-site I according to its crystal structure (PDB 3AGY). The mass density map is in cyan (meshes). EEVD peptide from the crystal structure is in red and the lowest energy poses are in green. **b** DNAJB1-site II according to its crystal structure (PDB 3AGY). EEVD peptide from the crystal structure is in red. The mass density map of the EEVD-peptide is in magenta (meshes) and the lowest structure is in green. **c** Sis1-site II (PDB 2B26) DNAJB1-site II according to its crystal structure (PDB 3AGY). EEVD peptide from the crystal structure is in red. The mass density map of the EEVD-peptide is in magenta (meshes) and the lowest structure is in green.

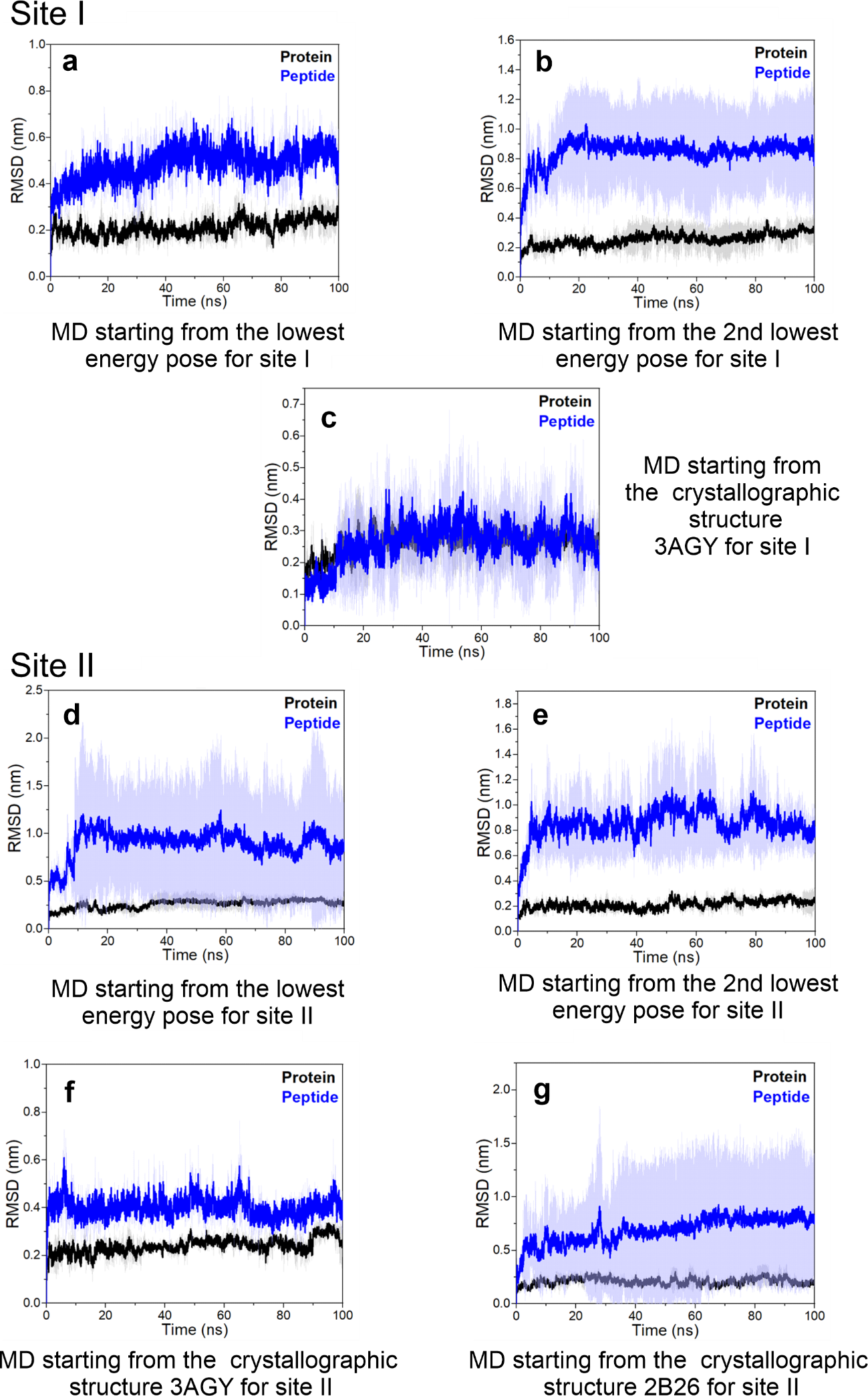

**Fig. S4 | Analysis of the parameters from 1.0 μs molecular dynamics simulations of the complex between CTDI sites I and II of JDPs and EEVD peptide.** **a, d** Values of root mean square deviations (RMSDs) of the backbone atoms of Sis1-CTDI from the lowest energy pose for site I and site II (black), respectively, and EEVD peptide (blue). **b, e** Values of RMSDs of the backbone atoms of Sis1-CTDI from the second lowest energy pose for site I and site II (black), respectively, and EEVD peptide (blue). **c, f** Values of RMSDs of the backbone atoms of DNAJB1-CTDI from the crystallographic structure (PDB 3AGY) for site I and site II (black), respectively, and EEVD peptide (blue). **g** Values of RMSDs of the backbone atoms of Sis1-CTDI from the crystallographic structure (PDB 2B26) for site II (black) and EEVD peptide (blue).

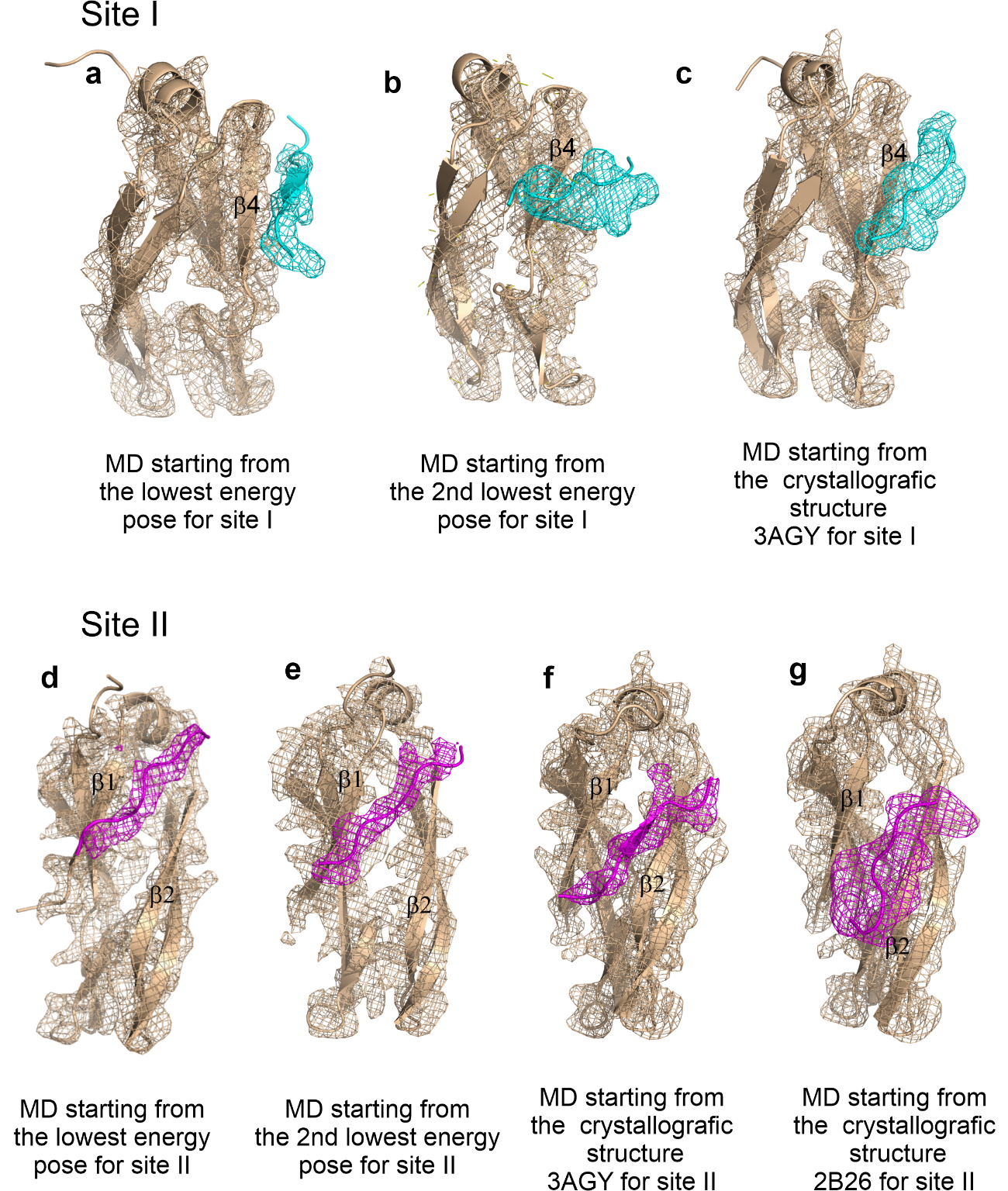

**Fig. S5 | Analysis of the stability between CTDI sites I and II of JDPs with EEVD peptide from 1.0 μs molecular dynamics (MD) simulations.** Cartoon representation of the CTDI and the mass density map (wheat color) from **a** the lowest energy pose for site I; **b** the second lowest energy pose for site I; **c** DNAJB1-site I according to its crystal structure (PDB 3AGY). The EEVD-peptide and its mass density map are in cyan (meshes); **d** the lowest energy pose for site II **e** the second lowest energy pose for site II; **f** DNAJB1-site II according to its crystal structure (PDB 3AGY); and **g** Sis1-site II according to its crystal structure (PDB 2B26). The EEVD-peptide and its mass density map are in magenta (meshes).

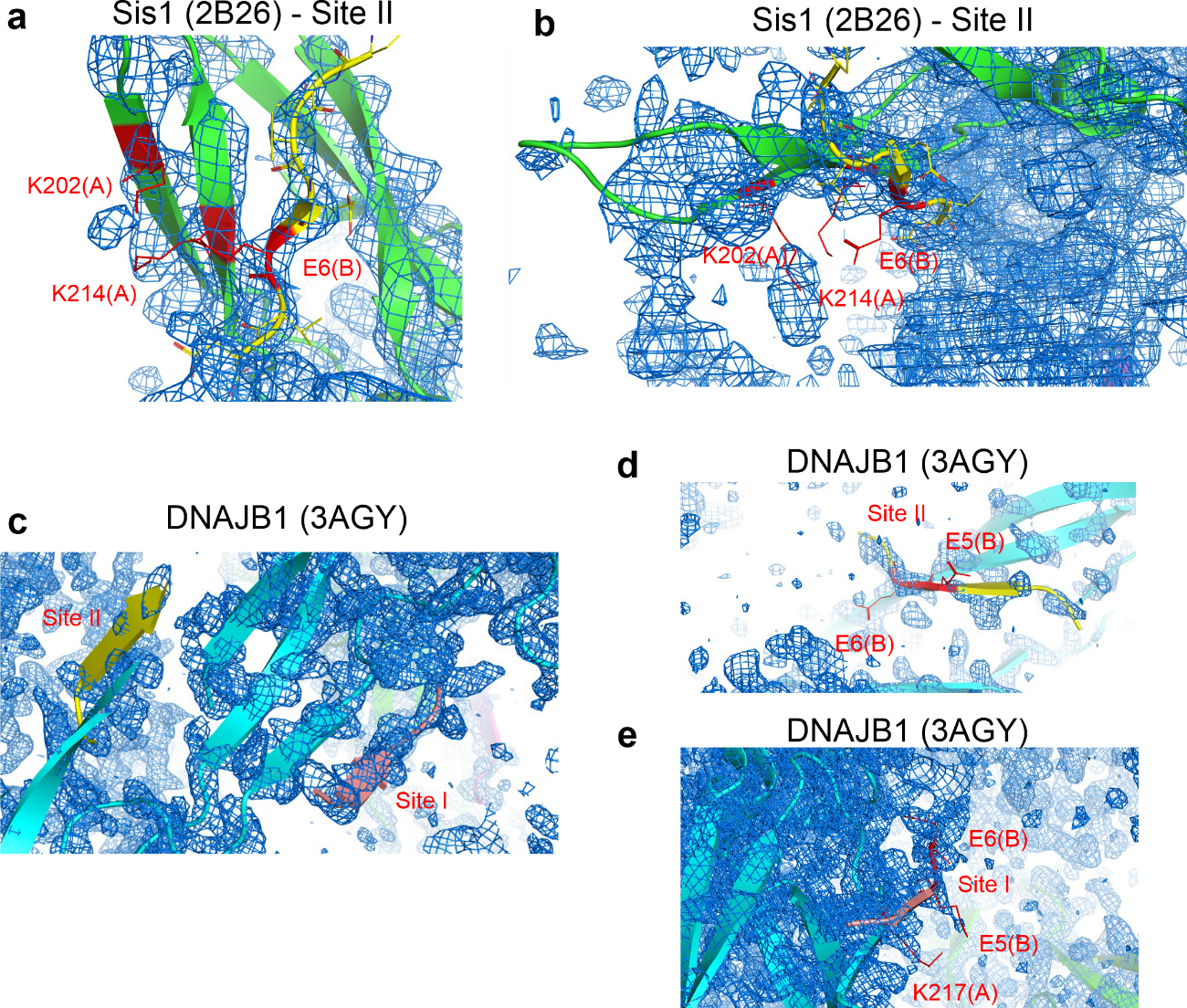

**Fig. S6 | Electron density map of crystal structures of JDPs bound to an EEVD peptide.** **a** and **b** Cartoon representation of CTDI of Sis1 (PDB 2B26) (green) with the EEVD peptide (yellow) bound at site II. The electron density map is in blue. Note that the EEVD backbone has a well-defined electron density, but side-chain information is lacking. **c-e** Cartoon representation of CTDI of DNAJB1 (PDB 3AGY) (cyan) with the EEVD peptide in yellow when bound to site II and in pink when bound to site I. The electron density map is in blue and lacks resolution at site II (**d)** but not at site I (**e)**, which is well-defined for residues P2, I4, E5 and E6.

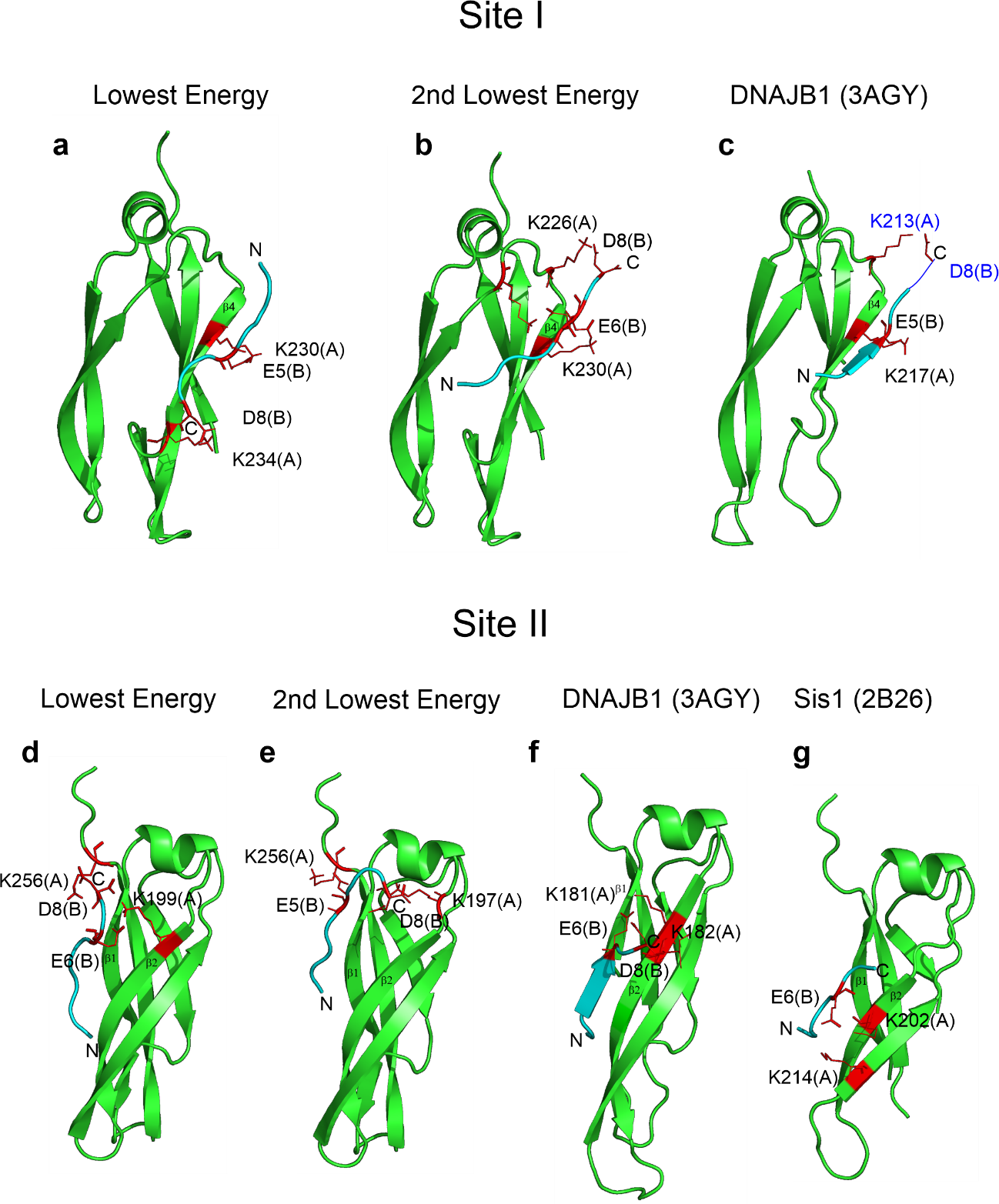

**Fig. S7. | Position and orientation of the EEVD-peptide at CTDI sites I and II of JDPs.** For site I: **a** The lowest energy structure from Sis1 (this work) shows that the EEVD-peptide has a parallel orientation relative to β4. **b** The second lowest structure diagram from Sis1 (this work) and **c** the crystal structure of DNAJB1 (PDB 3AGY) shows that the EEVD-peptide has an anti-parallel orientation relative to β4. Residues in dark blue are missing in the PDB structure and were drawn. For site II: **d** The lowest and **e** the second lowest structure are shown. The lowest energy structure from Sis1 (this work) interacts between β1 and β2 of CTDI. EEVD has the same anti-parallel orientation to β2 for **f** in the crystal DNAJB1 (PDB 3AGY) and **g** in the crystal Sis1 (PDB 2B26). CTDI structures are shown in green, EEVD peptide is labeled in cyan and salt bridges are shown in red.

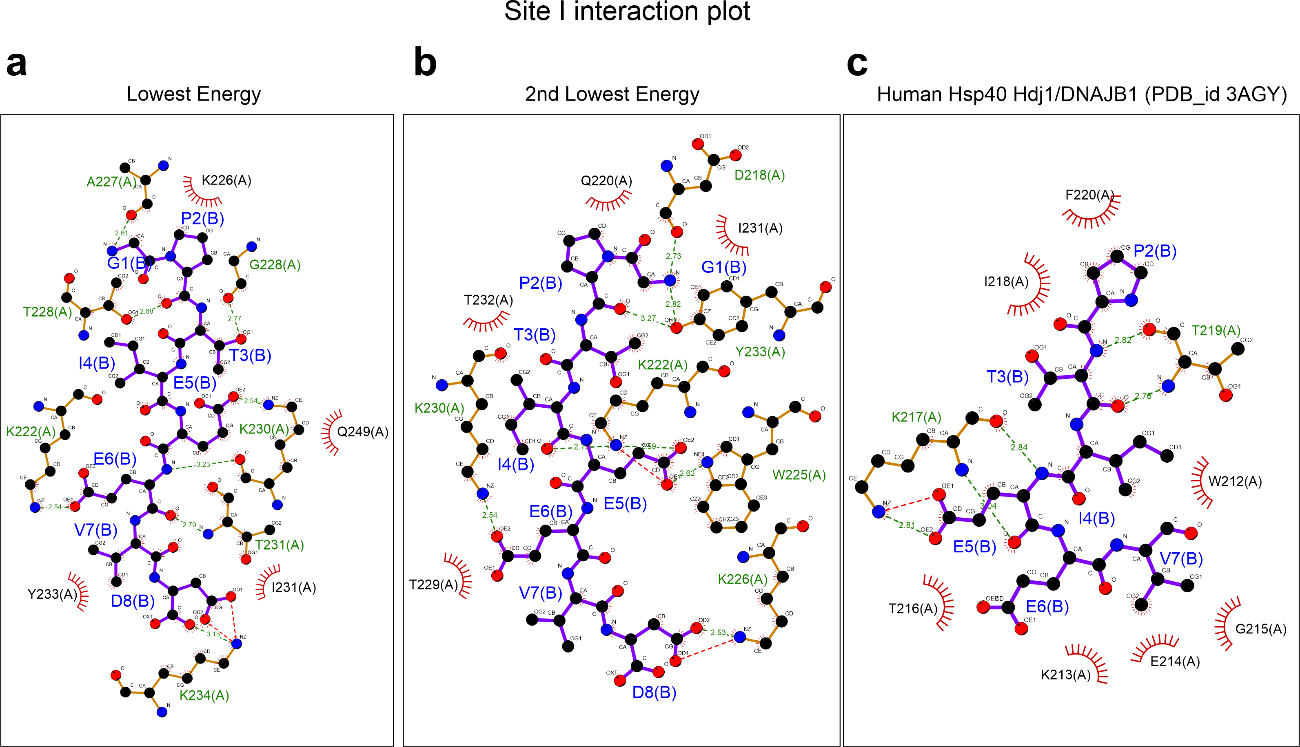

**Fig. S8. | EEVD-peptide interactions with the site I of JDPs CTDI. a** Lowest energy and **b** Second lowest energy diagram from Sis1 (this work). **c** Human DNAJB1 (PDB 3AGY). The plots were generated by Ligplot+ v.2.2.4. EEVD-peptide residues are shown in blue. Spoked arcs indicate that Sis1 residues are involved in hydrophobic contacts and green-labeled residues are involved in hydrogen bonds.

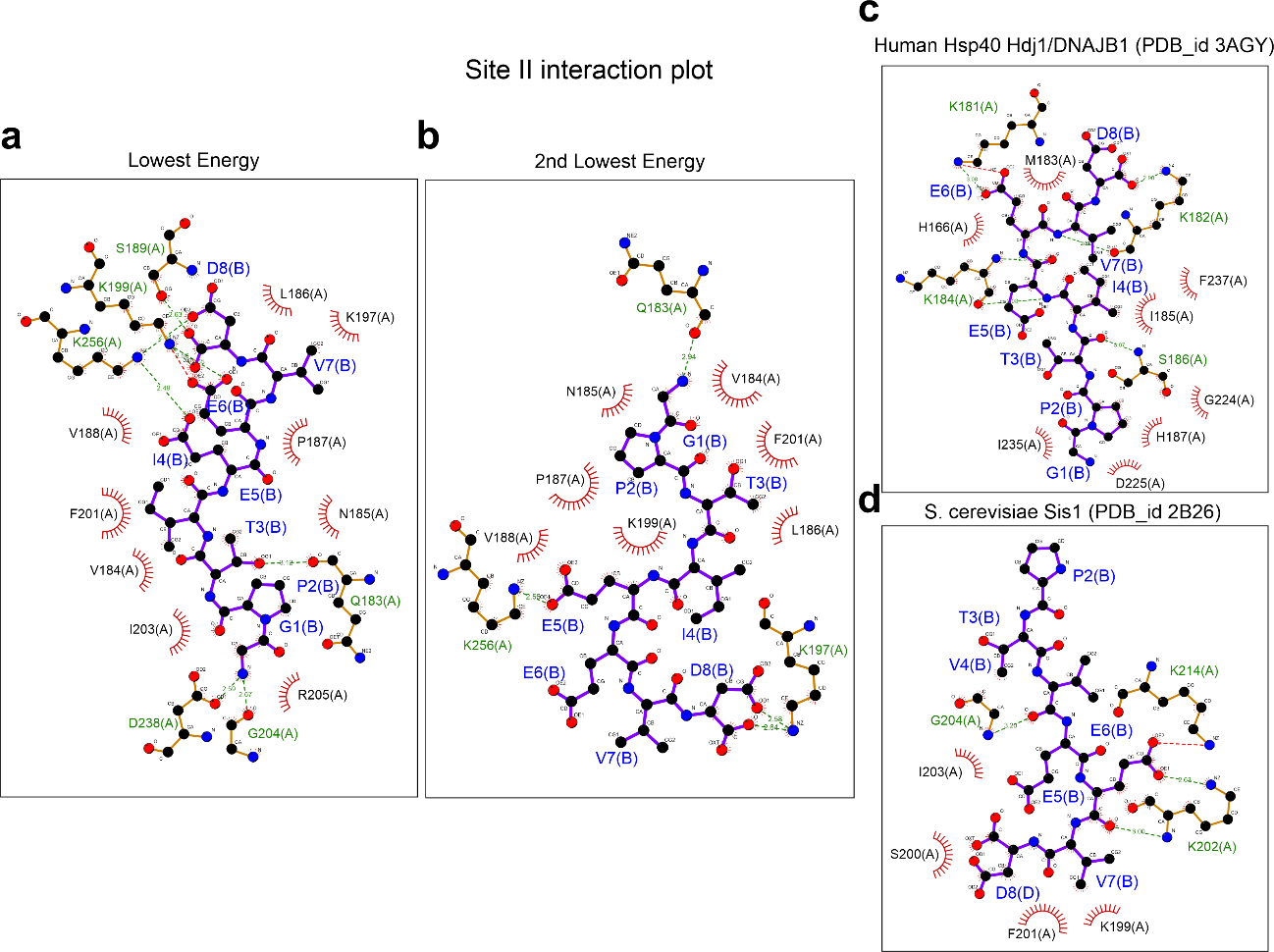

**Fig. S9. | EEVD-peptide interactions with the site II of JDPs CTDI. a** Lowest energy and **b** Second lowest energy diagram from Sis1 (this work). **c** Human DNAJB1 (PDB 3AGY). **d** Sis1 (data from PDB 2B26). The plots were generated by Ligplot+ v.2.2.4. EEVD-peptide residues are shown in blue. Spoked arcs indicate that Sis1 residues are involved in hydrophobic contacts and green-labeled residues are involved in hydrogen bonds.

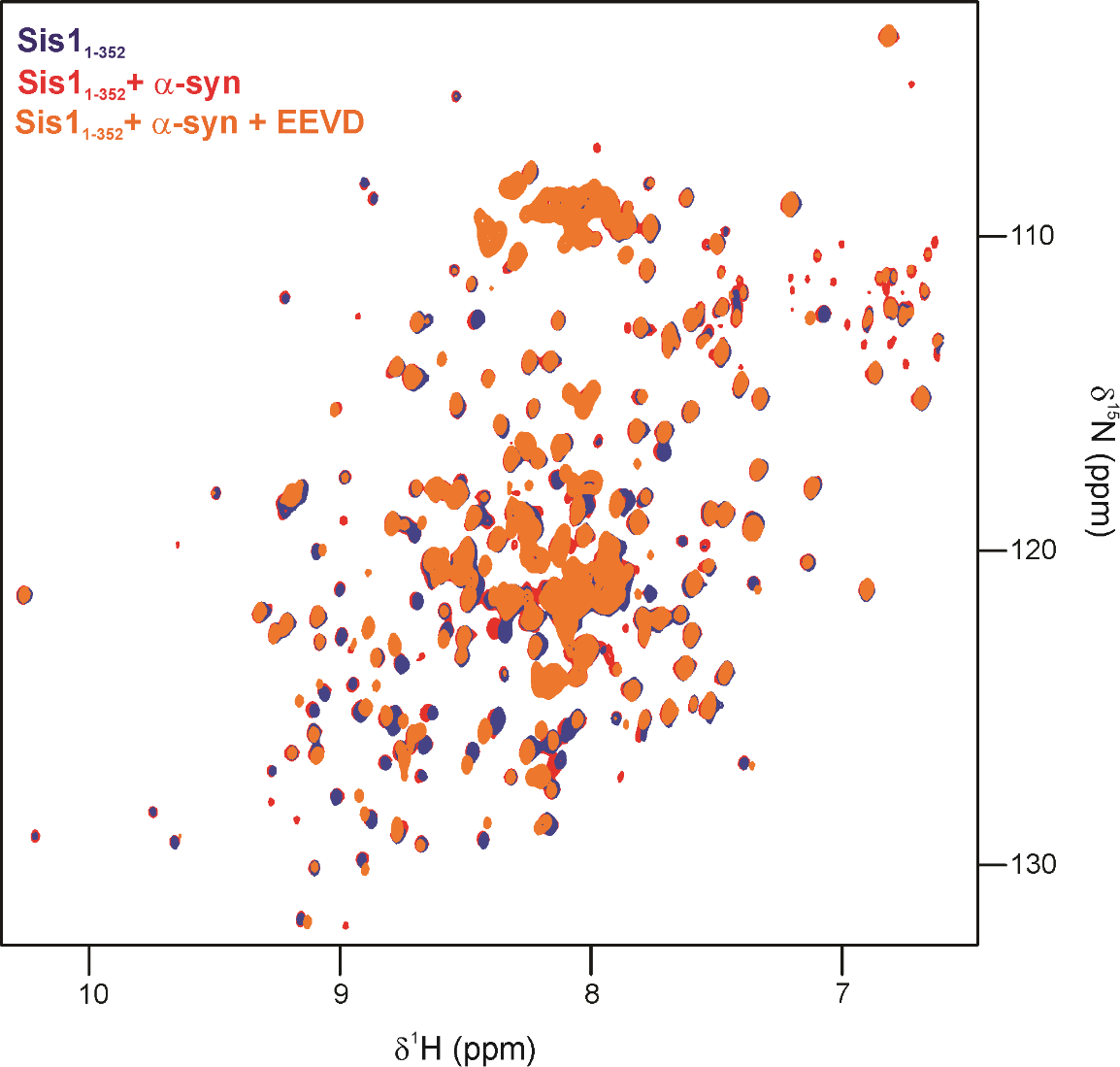

**Fig. S10 |** **Overlay of 2D ^1^H,^15^N TROSY HSQC spectra collected for ^2^H-^15^N-Sis1_1-352_ + α-syn and EEVD peptide.** Full spectra of Sis1_1-352_ free (blue), upon interaction with α-syn (red) and α-syn+ EEVD (orange).

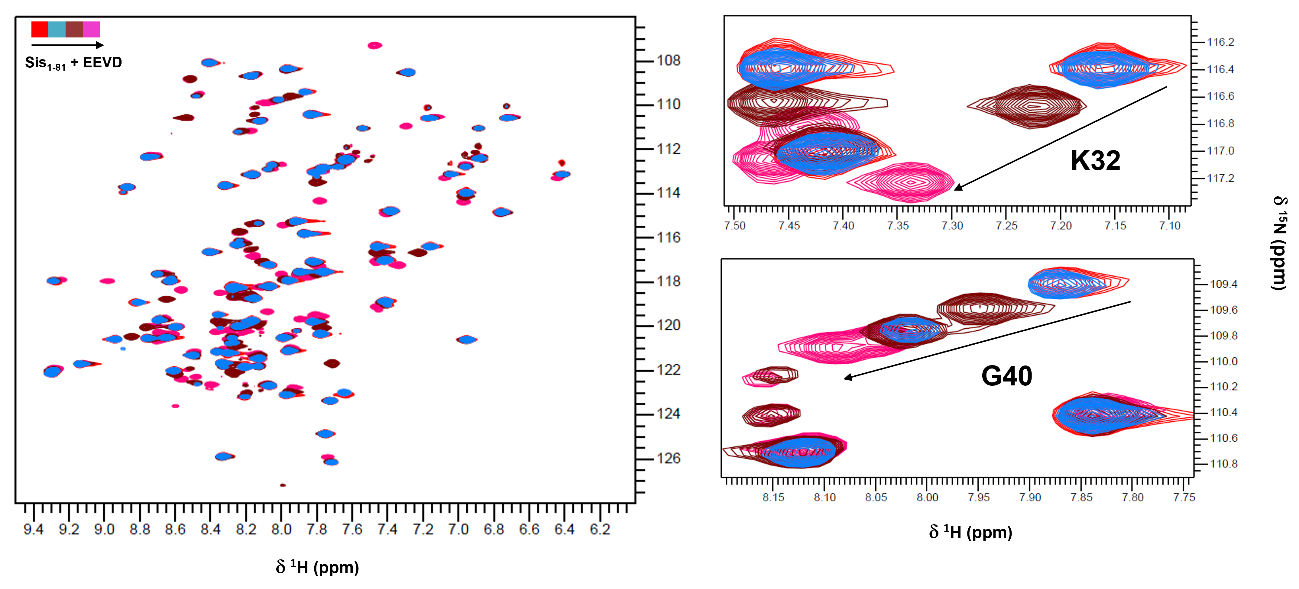
**Fig. S11 |** **Overlay of 2D ^1^H,^15^N HSQC spectra collected for ^15^N-Sis1_1-81_ at increasing EEVD-peptide concentration**. Sis1_1-81_ free (red), Sis1_1-81_ + 1 mM EEVD-peptide (blue), Sis1_1-81_ + 2 mM EEVD-peptide (brown) and Sis1_1-81_ + 4 mM EEVD-peptide (magenta). Left: full spectra; right: differential chemical shift data for residues K32 and G40 of Sis1_1-81_ which are involved in binding with the EEVD-peptide

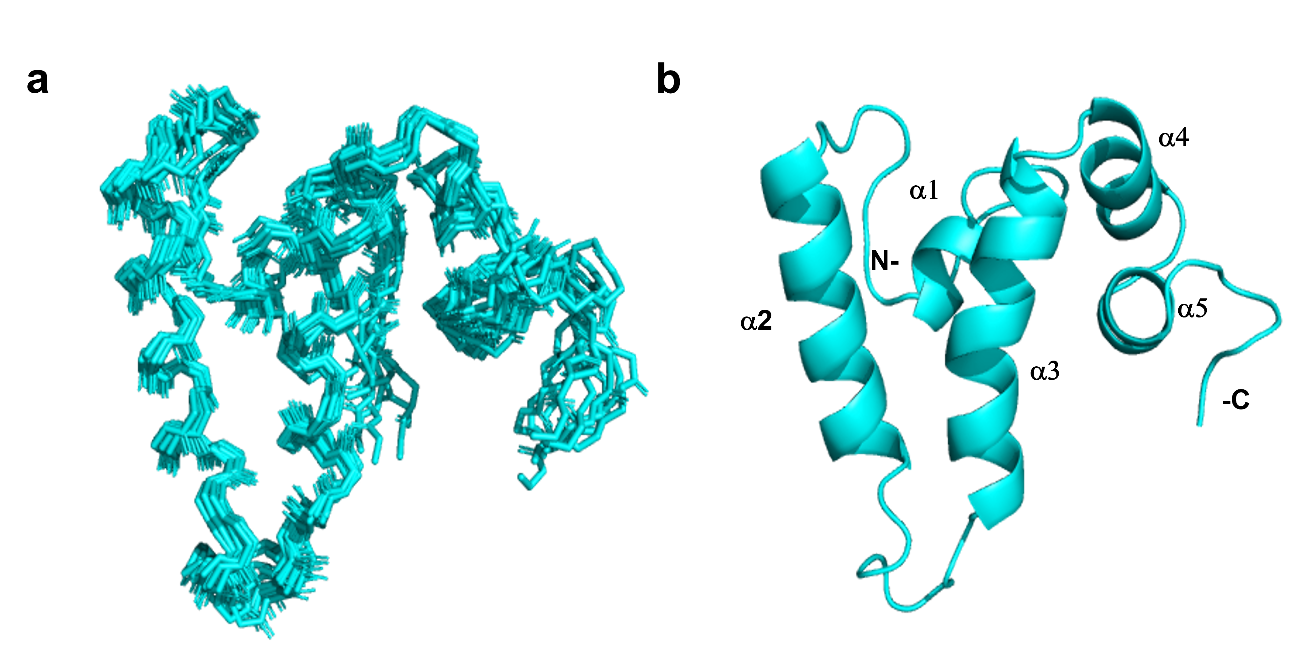

**Fig. S12 | Solution structure of Sis1_1-81_ bound to the EEVD-peptide a** Ensemble of 20 lowest-energy structures of the J-domain of Sis1 (1-81) with the HSP70 C-terminal peptide GPTIEEVD (Sis1_1-81_:EEVD-bound conformation). **b** Ribbon representation of the lowest-energy structure of the Sis1_1-81_:EEVD-bound conformation. N- and C-termini and α-helices (α1 to α5 are shown in the structure.

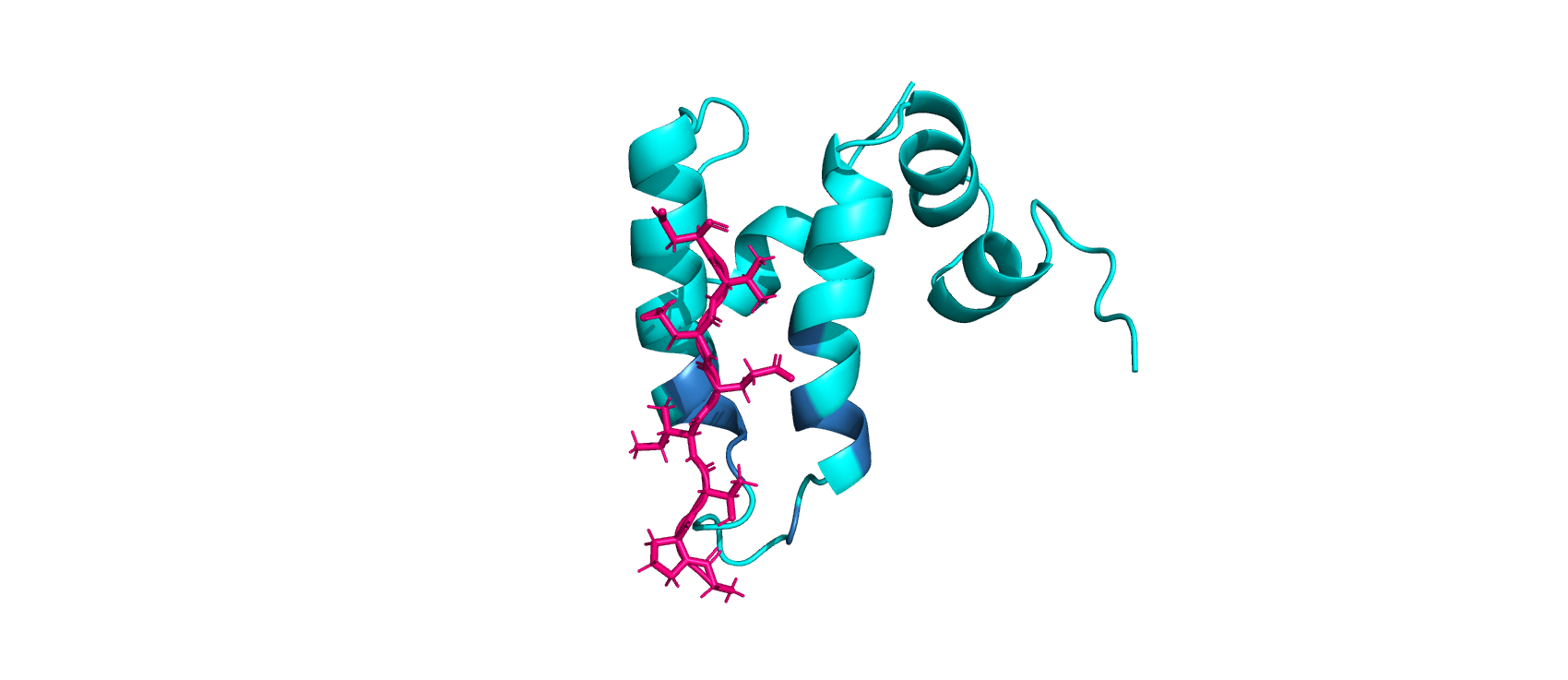

**Fig. S13 | Structure of the Sis1_1-81_:EEVD-peptide complex:** Ribbon representation of the lowest-energy structure of the Sis1_1-81_:EEVD-peptide complex calculated from HADDOCK using an NMR data-based model. The peptide extends between α2 and α3 with its N-terminal at the HPD-loop. Sis1_1-81_ is shown in cyan and the EEVD peptide is in pink. The structure is in good agreement with the CSP data (blue), PDB 8EOD.

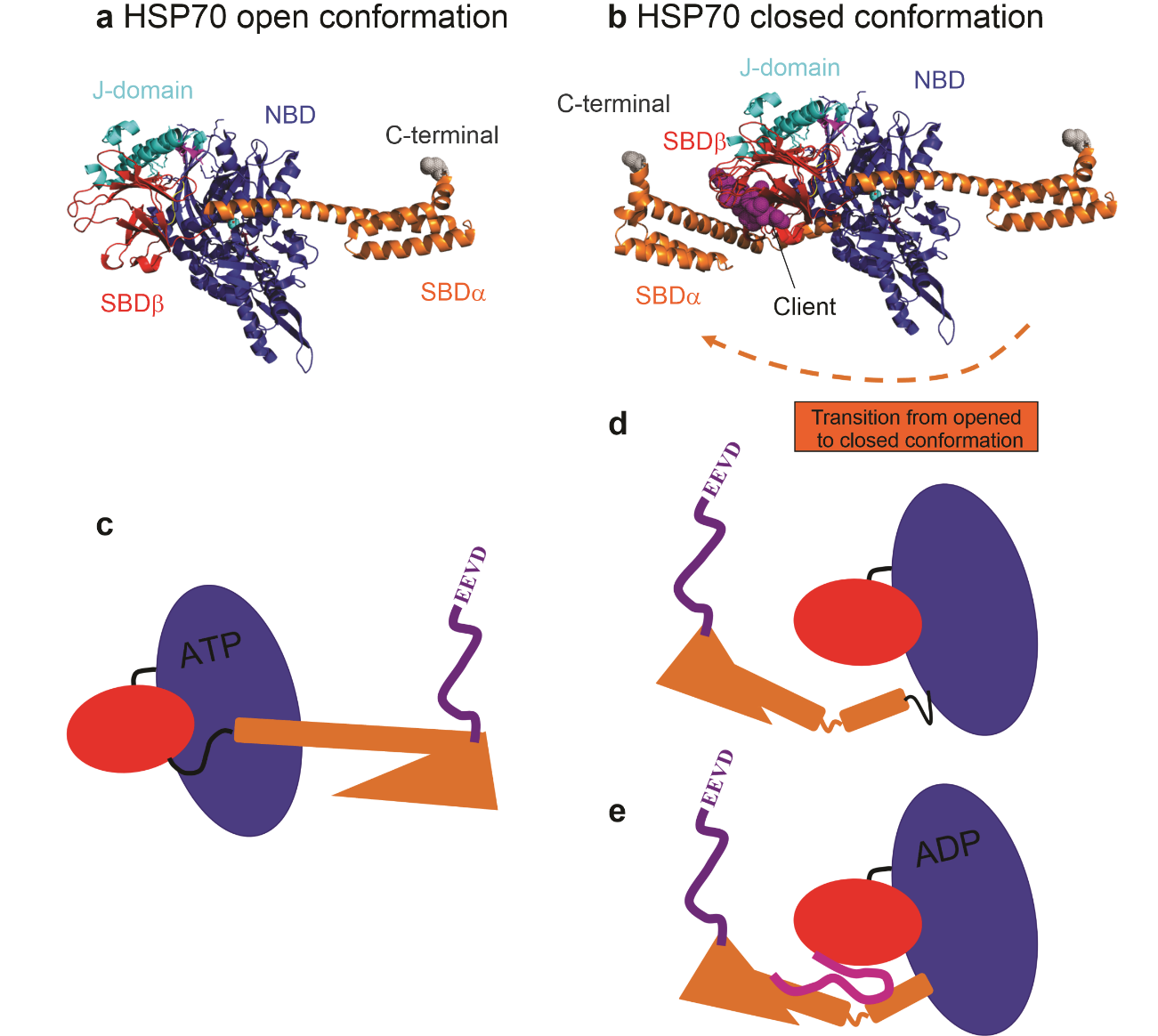

**Fig. S14 |** Structure and schematic drawing of the HSP70 open and closed conformations. Note that the schematic drawing replicates the position and orientation of each domain of HSP70: NBD (blue), SBDα (orange), SBDβ (red). **a** SBD open conformation based on the structure of the ATP-bound *E. coli* DNAK complexed with the J-domain (5NRO^2^). **b** Structure of the *E. coli* DNAK complexed with a client protein (1DKX^1^) superposed with the structure of *E. coli* DNAK complexed with the J-domain (5NRO^2^). **c**. Schematic view of the open conformational state. **d**. Schematic view of the closed conformational state in the apo state. The figure illustrates the conserved linker between the SBD and NBD. In this state the linker is exposed. **e.** Schematic view of the closed conformational state in the ADP-bound state. In this state the linker is inserted into a cavity of NBD. The linker is essential for the allosteric mechanism illustrated in Figure 7. The position and orientation of the C-terminal IDR containing the EEVD peptide are compatible with the Alphafold prediction (not shown). The Alphafold structure of the open conformation of SSA1, the yeast homologous of HSC70 and DNAK is significantly similar and superposable to the structure of DNAK (5NRO).

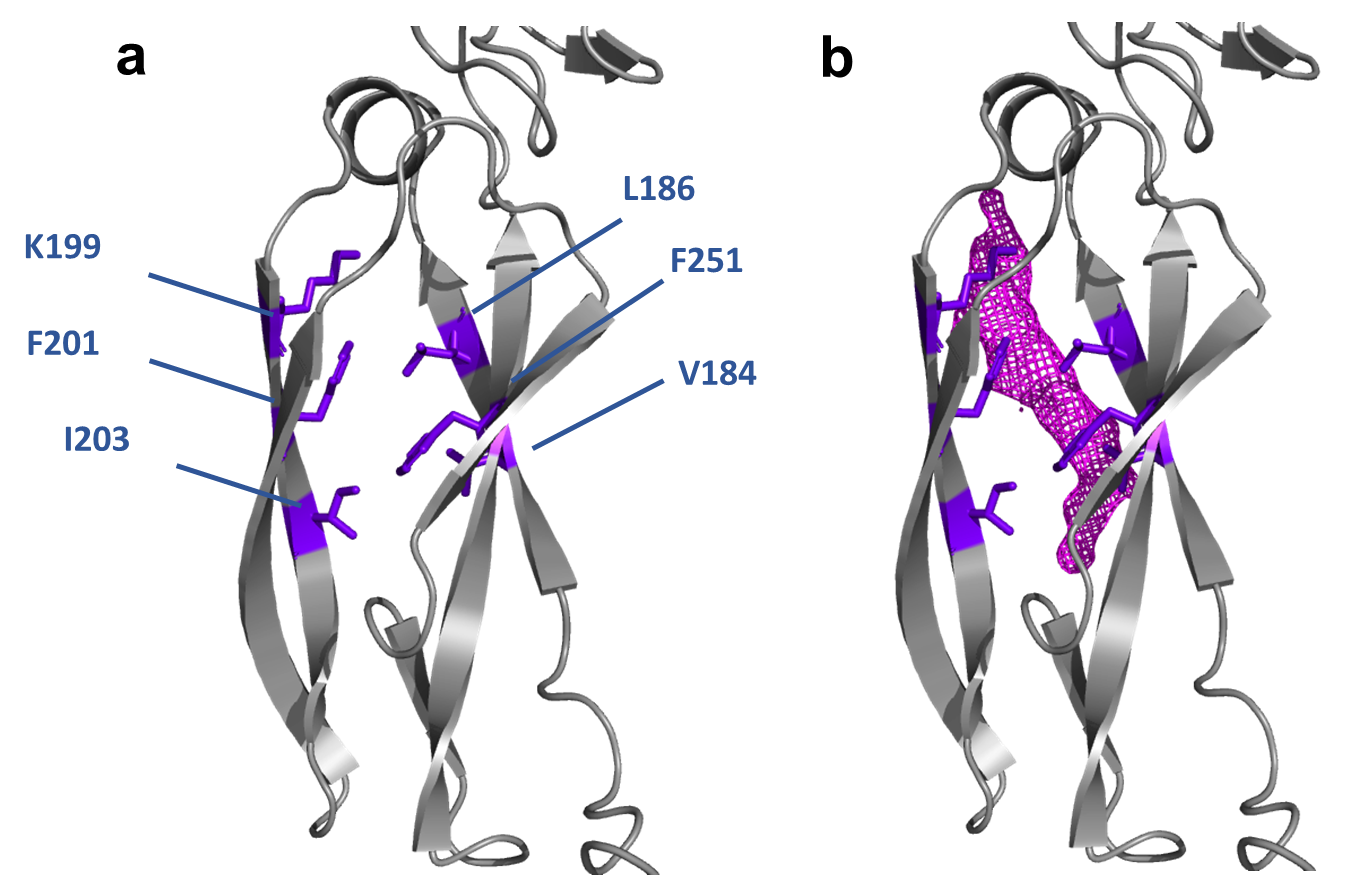

**Fig. S15 | CTDI peptide binding site of Sis1**. **a** Residues V184, L186, K199, F201, K203, and F251 (blue) are previously described to form a large hydrophobic depression that is likely involved in client protein binding. **b** The atomic probability density map of the EEVD-peptide on CTDI-Sis1 (magenta; this work) interacts with the same region described to be involved to bind to the substrate.

**Table S1**. NMR-derived restraints and structural statistics for the 20 best water-refined structures of Sis1_1-81_:EEVD-bound conformation.

| Number of experimental restraints | |
| --- | --- |
| Total NOE distance restraints (ambiguous) | 1206 |
| Total NOE distance restraints (unambiguous) | 2750 |
| Total dihedral angle restraints | 132 |
| Ambiguous | |
| Short range (\|I – j\| ≤ 1) | 835 |
| Medium range (2 ≤ \|I – j\| ≤ 4) | 248 |
| Long range (\|I – j\| > 4) | 123 |
| Unambiguous | |
| Short range (\|I – j\| ≤1) | 1821 |
| Medium range (2 ≤ \|I – j\| ≤ 4) | 560 |
| Long range (\|I – j\| > 4) | 369 |
| Restraints violations | |
| distance constraints (>0.3 Å) | 0 |
| dihedral angle constraints (>5°) | 0 |
| RMSD from average structure (Å) | |
| Backbone (4-76) | 0.8 |
| Backbone, all residues | 1.4 |
| Heavy atoms (4-76) | 1.2 |
| Heavy atoms, all residues | 1.7 |
| Ramachandran plot of ordered residues – Procheck (%) | |
| Most favored regions | 93.6 |
| Allowed regions | 6.4 |
| Generously allowed | 0 |
| Disallowed | 0 |
| Ramachandran plot of ordered residues – Molprobity (%) | |
| Most favored regions | 97.7 |
| Allowed regions | 2.2 |
| Disallowed | 0.1 |
| CNS Energy Score (kcal/mol) | |
| E_tot_ | -3490 ± 88 |
| E_bound_ | 12.3 ± 0.6 |
| E_angle_ | 68 ± 4 |
| E_impr_ | 128 ±15 |
| E_dihed_ | 378 ± 4 |
| E_vdw_ | -763 ± 10 |
| E_elec_ | -3313 ± 88 |

**Table S2:** Haddock results of molecular docking for Sis1_1-81_:EEVD complex

| **Parameters** | **Sis1_1-81_:EEVD complex** |
| --- | --- |
| HADDOCK score | -57.8 ± 6.5 |
| Cluster size | 7 |
| RMSD | 0.7 ± 0.5 |
| Van der Waals energy | -12 ± 11 |
| Electrostatic energy | -242 ± 48 |
| Desolvation energy | 0 ± 3 |
| Restraints violation energy | 29 ± 14 |
| Buried Surface Area | 832 ± 32 |
| Z-score | -2.4 |

**Table S3.** Hydrogen bonds between Sis_1-81_ and EEVD peptide with more than 10 % of persistence throughout the 1 μs MD simulation. The residues and atoms of the EEVD peptide are in red, while the residues and atoms of Sis_1-81_ are in black. Note that some of the hydrogen bonds participate in the intermolecular salt bridges that stabilize the Sis_1-81_:EEVD complex.

| **Donor** | **Atom** | **Acceptor** | **Atom** | **Prevalence %** | **Salt Bridge** |
| --- | --- | --- | --- | --- | --- |
| GLU5 | N | TYR26 | OH | 92.6 | - |
| ASP8 | N | ASN56 | OD1 | 82.3 | - |
| LYS23 | NZ | GLU6 | OE1 | 32.9 | yes |
| HIS34 | NE2 | PRO2 | O | 32.1 | - |
| LYS23 | NZ | GLU6 | OE2 | 30.2 | yes |
| ASN56 | ND2 | ASP8 | OC2 | 26.9 | - |
| ASN56 | ND2 | ASP8 | OC1 | 26.7 | - |
| ARG27 | NH2 | GLU6 | OE2 | 21.7 | yes |
| ARG27 | NH2 | GLU6 | OE1 | 21.4 | yes |

**Table S3 |** Distance restraints semi-quantitatively calibrated from the PRE data for CTDI-Sis1_1-352_:EEVD using for molecular docking

assign (segid "	A" and resid	187 and name	HN) (segid "	B" and resid	1 and name HN) 18.1 6.1 3 weight 1.000

assign (segid "	A" and resid	202 and name	HN) (segid "	B" and resid	1 and name HN) 26.8 14.8 3 weight 1.000

assign (segid "	A" and resid	206 and name	HN) (segid "	B" and resid	1 and name HN) 13.2 1.2 3 weight 1.000

assign (segid "	A" and resid	234 and name	HN) (segid "	B" and resid	1 and name HN) 26.9 14.9 3 weight 1.000

assign (segid "	A" and resid	240 and name	HN) (segid "	B" and resid	1 and name HN) 21.1 9.1 3 weight 1.000

assign (segid "	A" and resid	265 and name	HN) (segid "	B" and resid	1 and name HN) 22.5 10.5 3 weight 1.000

assign (segid "	A" and resid	286 and name	HN) (segid "	B" and resid	1 and name HN) 23.1 11.1 3 weight 1.000

assign (segid "	A" and resid	290 and name	HN) (segid "	B" and resid	1 and name HN) 25 13 3 weight 1.000

assign (segid "	A" and resid	331 and name	HN) (segid "	B" and resid	1 and name HN) 27.1 15.1 3 weight 1.000

assign (segid "	A" and resid	187 and name	HN) (segid "	B" and resid	1 and name N) 18.1 6.1 3 weight 1.000

assign (segid "	A" and resid	202 and name	HN) (segid "	B" and resid	1 and name N) 26.8 14.8 3 weight 1.000

assign (segid "	A" and resid	206 and name	HN) (segid "	B" and resid	1 and name N) 13.2 1.2 3 weight 1.000

assign (segid "	A" and resid	234 and name	HN) (segid "	B" and resid	1 and name N) 26.9 14.9 3 weight 1.000

assign (segid "	A" and resid	240 and name	HN) (segid "	B" and resid	1 and name N) 21.1 9.1 3 weight 1.000

assign (segid "	A" and resid	265 and name	HN) (segid "	B" and resid	1 and name N) 22.5 10.5 3 weight 1.000

assign (segid "	A" and resid	286 and name	HN) (segid "	B" and resid	1 and name N) 23.1 11.1 3 weight 1.000

assign (segid "	A" and resid	290 and name	HN) (segid "	B" and resid	1 and name N) 25 13 3 weight 1.000

assign (segid "	A" and resid	331 and name	HN) (segid "	B" and resid	1 and name N) 27.1 15.1 3 weight 1.000

assign (segid "	A" and resid	187 and name	HN) (segid "	B" and resid	1 and name CA) 18.1 6.1 3 weight 1.000

assign (segid "	A" and resid	202 and name	HN) (segid "	B" and resid	1 and name CA) 26.8 14.8 3 weight 1.000

assign (segid "	A" and resid	206 and name	HN) (segid "	B" and resid	1 and name CA) 13.2 1.2 3 weight 1.000

assign (segid "	A" and resid	234 and name	HN) (segid "	B" and resid	1 and name CA) 26.9 14.9 3 weight 1.000

assign (segid "	A" and resid	240 and name	HN) (segid "	B" and resid	1 and name CA) 21.1 9.1 3 weight 1.000

assign (segid "	A" and resid	265 and name	HN) (segid "	B" and resid	1 and name CA) 22.5 10.5 3 weight 1.000

assign (segid "	A" and resid	286 and name	HN) (segid "	B" and resid	1 and name CA) 23.1 11.1 3 weight 1.000

assign (segid "	A" and resid	290 and name	HN) (segid "	B" and resid	1 and name CA) 25 13 3 weight 1.000

assign (segid "	A" and resid	331 and name	HN) (segid "	B" and resid	1 and name CA) 27.1 15.1 3 weight 1.000

assign (segid "	A" and resid	187 and name	HN) (segid "	B" and resid	1 and name CB) 18.1 6.1 3 weight 1.000

assign (segid "	A" and resid	202 and name	HN) (segid "	B" and resid	1 and name CB) 26.8 14.8 3 weight 1.000

assign (segid "	A" and resid	206 and name	HN) (segid "	B" and resid	1 and name CB) 13.2 1.2 3 weight 1.000

assign (segid "	A" and resid	234 and name	HN) (segid "	B" and resid	1 and name CB) 26.9 14.9 3 weight 1.000

assign (segid "	A" and resid	240 and name	HN) (segid "	B" and resid	1 and name CB) 21.1 9.1 3 weight 1.000

assign (segid "	A" and resid	265 and name	HN) (segid "	B" and resid	1 and name CB) 22.5 10.5 3 weight 1.000

assign (segid "	A" and resid	286 and name	HN) (segid "	B" and resid	1 and name CB) 23.1 11.1 3 weight 1.000

assign (segid "	A" and resid	290 and name	HN) (segid "	B" and resid	1 and name CB) 25 13 3 weight 1.000

assign (segid "	A" and resid	331 and name	HN) (segid "	B" and resid	1 and name CB) 27.1 15.1 3 weight 1.000

assign (segid "	A" and resid	187 and name	HN) (segid "	B" and resid	1 and name C) 18.1 6.1 3 weight 1.000

assign (segid "	A" and resid	202 and name	HN) (segid "	B" and resid	1 and name C) 26.8 14.8 3 weight 1.000

assign (segid "	A" and resid	206 and name	HN) (segid "	B" and resid	1 and name C) 13.2 1.2 3 weight 1.000

assign (segid "	A" and resid	234 and name	HN) (segid "	B" and resid	1 and name C) 26.9 14.9 3 weight 1.000

assign (segid "	A" and resid	240 and name	HN) (segid "	B" and resid	1 and name C) 21.1 9.1 3 weight 1.000

assign (segid "	A" and resid	265 and name	HN) (segid "	B" and resid	1 and name C) 22.5 10.5 3 weight 1.000

assign (segid "	A" and resid	286 and name	HN) (segid "	B" and resid	1 and name C) 23.1 11.1 3 weight 1.000

assign (segid "	A" and resid	290 and name	HN) (segid "	B" and resid	1 and name C) 25 13 3 weight 1.000

assign (segid "	A" and resid	331 and name	HN) (segid "	B" and resid	1 and name C) 27.1 15.1 3 weight 1.000

assign (segid "	A" and resid	187 and name	HN) (segid "	B" and resid	1 and name HA) 18.1 6.1 3 weight 1.000

assign (segid "	A" and resid	202 and name	HN) (segid "	B" and resid	1 and name HA) 26.8 14.8 3 weight 1.000

assign (segid "	A" and resid	206 and name	HN) (segid "	B" and resid	1 and name HA) 13.2 1.2 3 weight 1.000

assign (segid "	A" and resid	234 and name	HN) (segid "	B" and resid	1 and name HA) 26.9 14.9 3 weight 1.000

assign (segid "	A" and resid	240 and name	HN) (segid "	B" and resid	1 and name HA) 21.1 9.1 3 weight 1.000

assign (segid "	A" and resid	265 and name	HN) (segid "	B" and resid	1 and name HA) 22.5 10.5 3 weight 1.000

assign (segid "	A" and resid	286 and name	HN) (segid "	B" and resid	1 and name HA) 23.1 11.1 3 weight 1.000

assign (segid "	A" and resid	290 and name	HN) (segid "	B" and resid	1 and name HA) 25 13 3 weight 1.000

assign (segid "	A" and resid	331 and name	HN) (segid "	B" and resid	1 and name HA) 27.1 15.1 3 weight 1.000

assign (segid "	A" and resid	187 and name	HN) (segid "	B" and resid	1 and name HB) 18.1 6.1 3 weight 1.000

assign (segid "	A" and resid	202 and name	HN) (segid "	B" and resid	1 and name HB) 26.8 14.8 3 weight 1.000

assign (segid "	A" and resid	206 and name	HN) (segid "	B" and resid	1 and name HB) 13.2 1.2 3 weight 1.000

assign (segid "	A" and resid	234 and name	HN) (segid "	B" and resid	1 and name HB) 26.9 14.9 3 weight 1.000

assign (segid "	A" and resid	240 and name	HN) (segid "	B" and resid	1 and name HB) 21.1 9.1 3 weight 1.000

assign (segid "	A" and resid	265 and name	HN) (segid "	B" and resid	1 and name HB) 22.5 10.5 3 weight 1.000

assign (segid "	A" and resid	286 and name	HN) (segid "	B" and resid	1 and name HB) 23.1 11.1 3 weight 1.000

assign (segid "	A" and resid	290 and name	HN) (segid "	B" and resid	1 and name HB) 25 13 3 weight 1.000

assign (segid "	A" and resid	331 and name	HN) (segid "	B" and resid	1 and name HB) 27.1 15.1 3 weight 1.000

**Table S4 |** Distance restraints semi-quantitatively calibrated from the PRE data for Sis1_1-81_:EEVD using for molecular docking

assign (segid "	A" and resid	10 and name	HN) (segid "	B" and resid	1 and name HN) 21 9 3 weight 1.000

assign (segid "	A" and resid	14 and name	HN) (segid "	B" and resid	1 and name HN) 26.7 14.7 3 weight 1.000

assign (segid "	A" and resid	24 and name	HN) (segid "	B" and resid	1 and name HN) 26.1 14.1 3 weight 1.000

assign (segid "	A" and resid	28 and name	HN) (segid "	B" and resid	1 and name HN) 17.4 5.4 3 weight 1.000

assign (segid "	A" and resid	30 and name	HN) (segid "	B" and resid	1 and name HN) 12 10.2 3 weight 1.000

assign (segid "	A" and resid	31 and name	HN) (segid "	B" and resid	1 and name HN) 16.2 4.2 3 weight 1.000

assign (segid "	A" and resid	32 and name	HN) (segid "	B" and resid	1 and name HN) 12.3 10.5 3 weight 1.000

assign (segid "	A" and resid	33 and name	HN) (segid "	B" and resid	1 and name HN) 14.3 12.5 3 weight 1.000

assign (segid "	A" and resid	34 and name	HN) (segid "	B" and resid	1 and name HN) 12.1 10.3 3 weight 1.000

assign (segid "	A" and resid	37 and name	HN) (segid "	B" and resid	1 and name HN) 31.3 19.3 3 weight 1.000

assign (segid "	A" and resid	40 and name	HN) (segid "	B" and resid	1 and name HN) 12.5 10.7 3 weight 1.000

assign (segid "	A" and resid	41 and name	HN) (segid "	B" and resid	1 and name HN) 17.8 5.8 3 weight 1.000

assign (segid "	A" and resid	43 and name	HN) (segid "	B" and resid	1 and name HN) 15.3 3.3 3 weight 1.000

assign (segid "	A" and resid	44 and name	HN) (segid "	B" and resid	1 and name HN) 16.2 4.2 3 weight 1.000

assign (segid "	A" and resid	45 and name	HN) (segid "	B" and resid	1 and name HN) 12 10.2 3 weight 1.000

assign (segid "	A" and resid	49 and name	HN) (segid "	B" and resid	1 and name HN) 12 10.2 3 weight 1.000

assign (segid "	A" and resid	54 and name	HN) (segid "	B" and resid	1 and name HN) 27.4 15.4 3 weight 1.000

assign (segid "	A" and resid	55 and name	HN) (segid "	B" and resid	1 and name HN) 29.5 17.5 3 weight 1.000

assign (segid "	A" and resid	10 and name	HN) (segid "	B" and resid	1 and name CA) 21 9 3 weight 1.000

assign (segid "	A" and resid	14 and name	HN) (segid "	B" and resid	1 and name CA) 26.7 14.7 3 weight 1.000

assign (segid "	A" and resid	24 and name	HN) (segid "	B" and resid	1 and name CA) 26.1 14.1 3 weight 1.000

assign (segid "	A" and resid	28 and name	HN) (segid "	B" and resid	1 and name CA) 17.4 5.4 3 weight 1.000

assign (segid "	A" and resid	30 and name	HN) (segid "	B" and resid	1 and name CA) 12 10.2 3 weight 1.000

assign (segid "	A" and resid	31 and name	HN) (segid "	B" and resid	1 and name CA) 16.2 4.2 3 weight 1.000

assign (segid "	A" and resid	32 and name	HN) (segid "	B" and resid	1 and name CA) 12.3 10.5 3 weight 1.000

assign (segid "	A" and resid	33 and name	HN) (segid "	B" and resid	1 and name CA) 14.3 12.5 3 weight 1.000

assign (segid "	A" and resid	34 and name	HN) (segid "	B" and resid	1 and name CA) 12.1 10.3 3 weight 1.000

assign (segid "	A" and resid	37 and name	HN) (segid "	B" and resid	1 and name CA) 31.3 19.3 3 weight 1.000

assign (segid "	A" and resid	40 and name	HN) (segid "	B" and resid	1 and name CA) 12.5 10.7 3 weight 1.000

assign (segid "	A" and resid	41 and name	HN) (segid "	B" and resid	1 and name CA) 17.8 5.8 3 weight 1.000

assign (segid "	A" and resid	43 and name	HN) (segid "	B" and resid	1 and name CA) 15.3 3.3 3 weight 1.000

assign (segid "	A" and resid	44 and name	HN) (segid "	B" and resid	1 and name CA) 16.2 4.2 3 weight 1.000

assign (segid "	A" and resid	45 and name	HN) (segid "	B" and resid	1 and name CA) 12 10.2 3 weight 1.000

assign (segid "	A" and resid	49 and name	HN) (segid "	B" and resid	1 and name CA) 12 10.2 3 weight 1.000

assign (segid "	A" and resid	54 and name	HN) (segid "	B" and resid	1 and name CA) 27.4 15.4 3 weight 1.000

assign (segid "	A" and resid	55 and name	HN) (segid "	B" and resid	1 and name CA) 29.5 17.5 3 weight 1.000

assign (segid "	A" and resid	10 and name	HN) (segid "	B" and resid	1 and name HA) 21 9 3 weight 1.000

assign (segid "	A" and resid	14 and name	HN) (segid "	B" and resid	1 and name HA) 26.7 14.7 3 weight 1.000

assign (segid "	A" and resid	24 and name	HN) (segid "	B" and resid	1 and name HA) 26.1 14.1 3 weight 1.000

assign (segid "	A" and resid	28 and name	HN) (segid "	B" and resid	1 and name HA) 17.4 5.4 3 weight 1.000

assign (segid "	A" and resid	30 and name	HN) (segid "	B" and resid	1 and name HA) 12 10.2 3 weight 1.000

assign (segid "	A" and resid	31 and name	HN) (segid "	B" and resid	1 and name HA) 16.2 4.2 3 weight 1.000

assign (segid "	A" and resid	32 and name	HN) (segid "	B" and resid	1 and name HA) 12.3 10.5 3 weight 1.000

assign (segid "	A" and resid	33 and name	HN) (segid "	B" and resid	1 and name HA) 14.3 12.5 3 weight 1.000

assign (segid "	A" and resid	34 and name	HN) (segid "	B" and resid	1 and name HA) 12.1 10.3 3 weight 1.000

assign (segid "	A" and resid	37 and name	HN) (segid "	B" and resid	1 and name HA) 31.3 19.3 3 weight 1.000

assign (segid "	A" and resid	40 and name	HN) (segid "	B" and resid	1 and name HA) 12.5 10.7 3 weight 1.000

assign (segid "	A" and resid	41 and name	HN) (segid "	B" and resid	1 and name HA) 17.8 5.8 3 weight 1.000

assign (segid "	A" and resid	43 and name	HN) (segid "	B" and resid	1 and name HA) 15.3 3.3 3 weight 1.000

assign (segid "	A" and resid	44 and name	HN) (segid "	B" and resid	1 and name HA) 16.2 4.2 3 weight 1.000

assign (segid "	A" and resid	45 and name	HN) (segid "	B" and resid	1 and name HA) 12 10.2 3 weight 1.000

assign (segid "	A" and resid	49 and name	HN) (segid "	B" and resid	1 and name HA) 12 10.2 3 weight 1.000

assign (segid "	A" and resid	54 and name	HN) (segid "	B" and resid	1 and name HA) 27.4 15.4 3 weight 1.000

assign (segid "	A" and resid	55 and name	HN) (segid "	B" and resid	1 and name HA) 29.5 17.5 3 weight 1.000

assign (segid "	A" and resid	10 and name	HN) (segid "	B" and resid	1 and name N) 21 9 3 weight 1.000

assign (segid "	A" and resid	14 and name	HN) (segid "	B" and resid	1 and name N) 26.7 14.7 3 weight 1.000

assign (segid "	A" and resid	24 and name	HN) (segid "	B" and resid	1 and name N) 26.1 14.1 3 weight 1.000

assign (segid "	A" and resid	28 and name	HN) (segid "	B" and resid	1 and name N) 17.4 5.4 3 weight 1.000

assign (segid "	A" and resid	30 and name	HN) (segid "	B" and resid	1 and name N) 12 10.2 3 weight 1.000

assign (segid "	A" and resid	31 and name	HN) (segid "	B" and resid	1 and name N) 16.2 4.2 3 weight 1.000

assign (segid "	A" and resid	32 and name	HN) (segid "	B" and resid	1 and name N) 12.3 10.5 3 weight 1.000

assign (segid "	A" and resid	33 and name	HN) (segid "	B" and resid	1 and name N) 14.3 12.5 3 weight 1.000

assign (segid "	A" and resid	34 and name	HN) (segid "	B" and resid	1 and name N) 12.1 10.3 3 weight 1.000

assign (segid "	A" and resid	37 and name	HN) (segid "	B" and resid	1 and name N) 31.3 19.3 3 weight 1.000

assign (segid "	A" and resid	40 and name	HN) (segid "	B" and resid	1 and name N) 12.5 10.7 3 weight 1.000

assign (segid "	A" and resid	41 and name	HN) (segid "	B" and resid	1 and name N) 17.8 5.8 3 weight 1.000

assign (segid "	A" and resid	43 and name	HN) (segid "	B" and resid	1 and name N) 15.3 3.3 3 weight 1.000

assign (segid "	A" and resid	44 and name	HN) (segid "	B" and resid	1 and name N) 16.2 4.2 3 weight 1.000

assign (segid "	A" and resid	45 and name	HN) (segid "	B" and resid	1 and name N) 12 10.2 3 weight 1.000

assign (segid "	A" and resid	49 and name	HN) (segid "	B" and resid	1 and name N) 12 10.2 3 weight 1.000

assign (segid "	A" and resid	54 and name	HN) (segid "	B" and resid	1 and name N) 27.4 15.4 3 weight 1.000

assign (segid "	A" and resid	55 and name	HN) (segid "	B" and resid	1 and name N) 29.5 17.5 3 weight 1.000

assign (segid "	A" and resid	10 and name	HN) (segid "	B" and resid	1 and name C) 21 9 3 weight 1.000

assign (segid "	A" and resid	14 and name	HN) (segid "	B" and resid	1 and name C) 26.7 14.7 3 weight 1.000

assign (segid "	A" and resid	24 and name	HN) (segid "	B" and resid	1 and name C) 26.1 14.1 3 weight 1.000

assign (segid "	A" and resid	28 and name	HN) (segid "	B" and resid	1 and name C) 17.4 5.4 3 weight 1.000

assign (segid "	A" and resid	30 and name	HN) (segid "	B" and resid	1 and name C) 12 10.2 3 weight 1.000

assign (segid "	A" and resid	31 and name	HN) (segid "	B" and resid	1 and name C) 16.2 4.2 3 weight 1.000

assign (segid "	A" and resid	32 and name	HN) (segid "	B" and resid	1 and name C) 12.3 10.5 3 weight 1.000

assign (segid "	A" and resid	33 and name	HN) (segid "	B" and resid	1 and name C) 14.3 12.5 3 weight 1.000

assign (segid "	A" and resid	34 and name	HN) (segid "	B" and resid	1 and name C) 12.1 10.3 3 weight 1.000

assign (segid "	A" and resid	37 and name	HN) (segid "	B" and resid	1 and name C) 31.3 19.3 3 weight 1.000

assign (segid "	A" and resid	40 and name	HN) (segid "	B" and resid	1 and name C) 12.5 10.7 3 weight 1.000

assign (segid "	A" and resid	41 and name	HN) (segid "	B" and resid	1 and name C) 17.8 5.8 3 weight 1.000

assign (segid "	A" and resid	43 and name	HN) (segid "	B" and resid	1 and name C) 15.3 3.3 3 weight 1.000

assign (segid "	A" and resid	44 and name	HN) (segid "	B" and resid	1 and name C) 16.2 4.2 3 weight 1.000

assign (segid "	A" and resid	45 and name	HN) (segid "	B" and resid	1 and name C) 12 10.2 3 weight 1.000

assign (segid "	A" and resid	49 and name	HN) (segid "	B" and resid	1 and name C) 12 10.2 3 weight 1.000

assign (segid "	A" and resid	54 and name	HN) (segid "	B" and resid	1 and name C) 27.4 15.4 3 weight 1.000

assign (segid "	A" and resid	55 and name	HN) (segid "	B" and resid	1 and name C) 29.5 17.5 3 weight 1.000

assign (segid "	A" and resid	10 and name	HN) (segid "	B" and resid	1 and name HB) 21 9 3 weight 1.000

assign (segid "	A" and resid	14 and name	HN) (segid "	B" and resid	1 and name HB) 26.7 14.7 3 weight 1.000

assign (segid "	A" and resid	24 and name	HN) (segid "	B" and resid	1 and name HB) 26.1 14.1 3 weight 1.000

assign (segid "	A" and resid	28 and name	HN) (segid "	B" and resid	1 and name HB) 17.4 5.4 3 weight 1.000

assign (segid "	A" and resid	30 and name	HN) (segid "	B" and resid	1 and name HB) 12 10.2 3 weight 1.000

assign (segid "	A" and resid	31 and name	HN) (segid "	B" and resid	1 and name HB) 16.2 4.2 3 weight 1.000

assign (segid "	A" and resid	32 and name	HN) (segid "	B" and resid	1 and name HB) 12.3 10.5 3 weight 1.000

assign (segid "	A" and resid	33 and name	HN) (segid "	B" and resid	1 and name HB) 14.3 12.5 3 weight 1.000

assign (segid "	A" and resid	34 and name	HN) (segid "	B" and resid	1 and name HB) 12.1 10.3 3 weight 1.000

assign (segid "	A" and resid	37 and name	HN) (segid "	B" and resid	1 and name HB) 31.3 19.3 3 weight 1.000

assign (segid "	A" and resid	40 and name	HN) (segid "	B" and resid	1 and name HB) 12.5 10.7 3 weight 1.000

assign (segid "	A" and resid	41 and name	HN) (segid "	B" and resid	1 and name HB) 17.8 5.8 3 weight 1.000

assign (segid "	A" and resid	43 and name	HN) (segid "	B" and resid	1 and name HB) 15.3 3.3 3 weight 1.000

assign (segid "	A" and resid	44 and name	HN) (segid "	B" and resid	1 and name HB) 16.2 4.2 3 weight 1.000

assign (segid "	A" and resid	45 and name	HN) (segid "	B" and resid	1 and name HB) 12 10.2 3 weight 1.000

assign (segid "	A" and resid	49 and name	HN) (segid "	B" and resid	1 and name HB) 12 10.2 3 weight 1.000

assign (segid "	A" and resid	54 and name	HN) (segid "	B" and resid	1 and name HB) 27.4 15.4 3 weight 1.000

assign (segid "	A" and resid	55 and name	HN) (segid "	B" and resid	1 and name HB) 29.5 17.5 3 weight 1.000

assign (segid "	A" and resid	10 and name	HN) (segid "	B" and resid	1 and name CB) 21 9 3 weight 1.000

assign (segid "	A" and resid	14 and name	HN) (segid "	B" and resid	1 and name CB) 26.7 14.7 3 weight 1.000

assign (segid "	A" and resid	24 and name	HN) (segid "	B" and resid	1 and name CB) 26.1 14.1 3 weight 1.000

assign (segid "	A" and resid	28 and name	HN) (segid "	B" and resid	1 and name CB) 17.4 5.4 3 weight 1.000

assign (segid "	A" and resid	30 and name	HN) (segid "	B" and resid	1 and name CB) 12 10.2 3 weight 1.000

assign (segid "	A" and resid	31 and name	HN) (segid "	B" and resid	1 and name CB) 16.2 4.2 3 weight 1.000

assign (segid "	A" and resid	32 and name	HN) (segid "	B" and resid	1 and name CB) 12.3 10.5 3 weight 1.000

assign (segid "	A" and resid	33 and name	HN) (segid "	B" and resid	1 and name CB) 14.3 12.5 3 weight 1.000

assign (segid "	A" and resid	34 and name	HN) (segid "	B" and resid	1 and name CB) 12.1 10.3 3 weight 1.000

assign (segid "	A" and resid	37 and name	HN) (segid "	B" and resid	1 and name CB) 31.3 19.3 3 weight 1.000

assign (segid "	A" and resid	40 and name	HN) (segid "	B" and resid	1 and name CB) 12.5 10.7 3 weight 1.000

assign (segid "	A" and resid	41 and name	HN) (segid "	B" and resid	1 and name CB) 17.8 5.8 3 weight 1.000

assign (segid "	A" and resid	43 and name	HN) (segid "	B" and resid	1 and name CB) 15.3 3.3 3 weight 1.000

assign (segid "	A" and resid	44 and name	HN) (segid "	B" and resid	1 and name CB) 16.2 4.2 3 weight 1.000

assign (segid "	A" and resid	45 and name	HN) (segid "	B" and resid	1 and name CB) 12 10.2 3 weight 1.000

assign (segid "	A" and resid	49 and name	HN) (segid "	B" and resid	1 and name CB) 12 10.2 3 weight 1.000

assign (segid "	A" and resid	54 and name	HN) (segid "	B" and resid	1 and name CB) 27.4 15.4 3 weight 1.000

assign (segid "	A" and resid	55 and name	HN) (segid "	B" and resid	1 and name CB) 29.5 17.5 3 weight 1.000
